# Stable and dynamic representations of value in the prefrontal cortex

**DOI:** 10.1101/794685

**Authors:** Pierre Enel, Joni Wallis, Erin Rich

## Abstract

The ability to associate positive and negative outcomes with predictive stimuli allows us to make optimal decisions. These stimulus-value associations are kept up to date by comparing an expected value with the experienced outcome. When a stimulus and its outcome are separated by a delay, the value associated with the stimulus must be held in mind for such comparisons to be possible, however little is known about the neural mechanisms that hold value representations online across delays. Temporarily remembering task-relevant information has been extensively studied in the context of item-specific working memory, and different hypotheses have suggested this ability requires either persistent or transient neuronal activity, with stable or dynamic representations respectively. To test these different hypotheses in the context of value representations, we recorded the spiking activity of neurons in the orbitofrontal and anterior cingulate cortex of two monkeys performing a task in which visual cues predicted a reward delivered after a short delay. We found that features of all hypotheses were simultaneously present in prefrontal activity and therefore no single hypothesis was exclusively supported. Instead, we report mixed dynamics that support robust, time invariant value representations while also encoding the information in a temporally specific manner. We suggest that this hybrid coding is important for optimal behavior and might be a critical mechanism supporting flexible cognitive abilities.

## Intro

Theories of value-based decision-making suggest that the brain computes values for different choice options in order to compare items with different qualities on a common scale [Padoa-Schioppa2011]. This attribution of a value relies on the association between an option and its outcome, and is learned by comparing the expected value of an option to the actual outcome experienced with it. However, when the presentation of an option is separated in time from its outcome, expected values must be temporarily held in memory for such comparisons to be possible.

The prefrontal cortex (PFC) plays a key role in temporarily maintaining and manipulating information across such temporal gaps, a process referred to as working memory (WM). While the lateral PFC is particularly involved in WM relating to cognitive information relevant to an internal goal [Constantinidis and Klingberg2016], it is less clear how value expectations are maintained in WM. Under the content model of WM, different prefrontal regions may be involved in maintaining and manipulating the type of information that they are specialized to process [Goldman-Rakic1996, Lara et al.2009]. From this view, maintaining an accurate estimation of expected values may rely more on those regions of PFC involved in learning and decision-making, such as the orbitofrontal cortex (OFC) or anterior cingulate cortex (ACC). The activity of single neurons in the OFC and ACC is known to reflect expected values associated with stimuli or actions [Rushworth and Behrens2008, Kennerley et al.2009, Amiez et al.2006], but little is know about the dynamics in either region that could bridge delays between outcomes and their predictive cues.

Here, we aimed to assess how value information is maintained across task delays in OFC and ACC. Neural mechanisms that maintain other domains of information in WM have been widely studied, but remain a subject of debate. Recently, this debate has focused on the nature of WM representations, culminating in two opposing theories [Constantinidis et al.2018, Lundqvist et al.2018]. One suggests that persistent and stable neuronal activity maintains WM representations across time [Wang2001, Riley and Constantinidis2016, Constantinidis et al.2018]. This hypothesis stems in part from the observation that neurons in the PFC maintain consistent activity patterns while monkeys wait during a delay to execute an action [Funahashi1989, Fuster1973]. These empirical results led to the development of well-studied neural network models in which attractor dynamics maintain information stably across time [Amit1995, Compte et al.2000].

However, more recently, the persistent activity hypothesis has been contested on several points. In various WM tasks, very little persistent activity is found in the lPFC of macaque monkeys [Stokes et al.2013, Lundqvist et al.2016, Rainer and Miller2002]. Nonetheless, the memoranda can be decoded throughout the delay from an evolving encoding pattern [Barak et al.2010], which prompted the development of a second theory, suggesting that WM involves dynamic representations and activity-silent synaptic encoding [Stokes et al.2017, Miller et al.2018]. Like the persistent activity hypothesis, this theory is also anchored in a theoretical framework, stemming from random recurrent neural networks of the reservoir computing type [Jaeger2001, Maass et al.2002]. Another theory postulates that WM could be implemented by the sequential activation of single neurons passing on information to bridge a delay [Rajan et al.2015, Harvey et al.2012]. While this idea is closer to the dynamic hypothesis than the stable/persistent hypothesis, unlike it, it predicts a well-defined pattern of activity at the level of single neurons, i.e. reliable sequences of activity.

Given these multiple perspectives, we investigated the nature of value representation in the OFC and ACC in a value-based decision making task in which monkeys were presented a reward predicting cue associated with a reward delivered after a delay. Similar to previous reports [Kennerley and Wallis2009], value encoding was found in both OFC and ACC during the delay. However, attempts at categorizing activity as persistent or transient, stable or dynamic, failed to exclusively support either view. Instead, both characteristics could be found in single unit and population activities. We demonstrate that it is possible to extract a purely stable or dynamic representation of value using different methods designed for detecting each.

These results support a more nuanced interpretation of OFC and ACC dynamics than any single hypothesis. Value representations in these cortical regions appear highly complex, as they include features of all theoretical frameworks. We hypothesize that these mixed dynamics occurring in the same neural population serve unique purposes. Stable representations allow downstream regions to read out memoranda with the same decoding scheme irrespective of the delay duration, while dynamic representations encode temporal information necessary to anticipate events and prepare behaviors. Such dynamics might allow the network to use a single value signal for different task-related purposes. From this view, a rich mixed dynamical regime could supply underlying mechanisms that support flexible cognitive abilities.

## Results

### Behavior

Two monkeys (subjects M and N) performed a value-based decision-making task in which visual stimuli were associated with juice rewards (Fig. 1A). Images predicted an amount and type of reward, which was either primary in the form of juice at the end of the trial, or secondary as a bar that increased proportionally to the value associated with the stimulus (Fig. 1B). The bar, always present on the screen, represented an amount of juice that was received when it cashed in every four trials. A total of 8 pictures comprised the set of stimuli associated with reward, with 4 different values matched between primary (juice) and secondary (bar) rewards in terms of the monkeys’ preferences [Rich and Wallis2016]. We will refer to reward value as an ordinal, from 1 (low value) to 4 (high value) for convenience. After the presentation of the pictures, monkeys were required to move a joystick either left or right depending on another visual cue in order to obtain the predicted reward. Joystick direction was unrelated to the reward-predicting picture, but if the incorrect response was made no reward was delivered.

**Figure 1:**
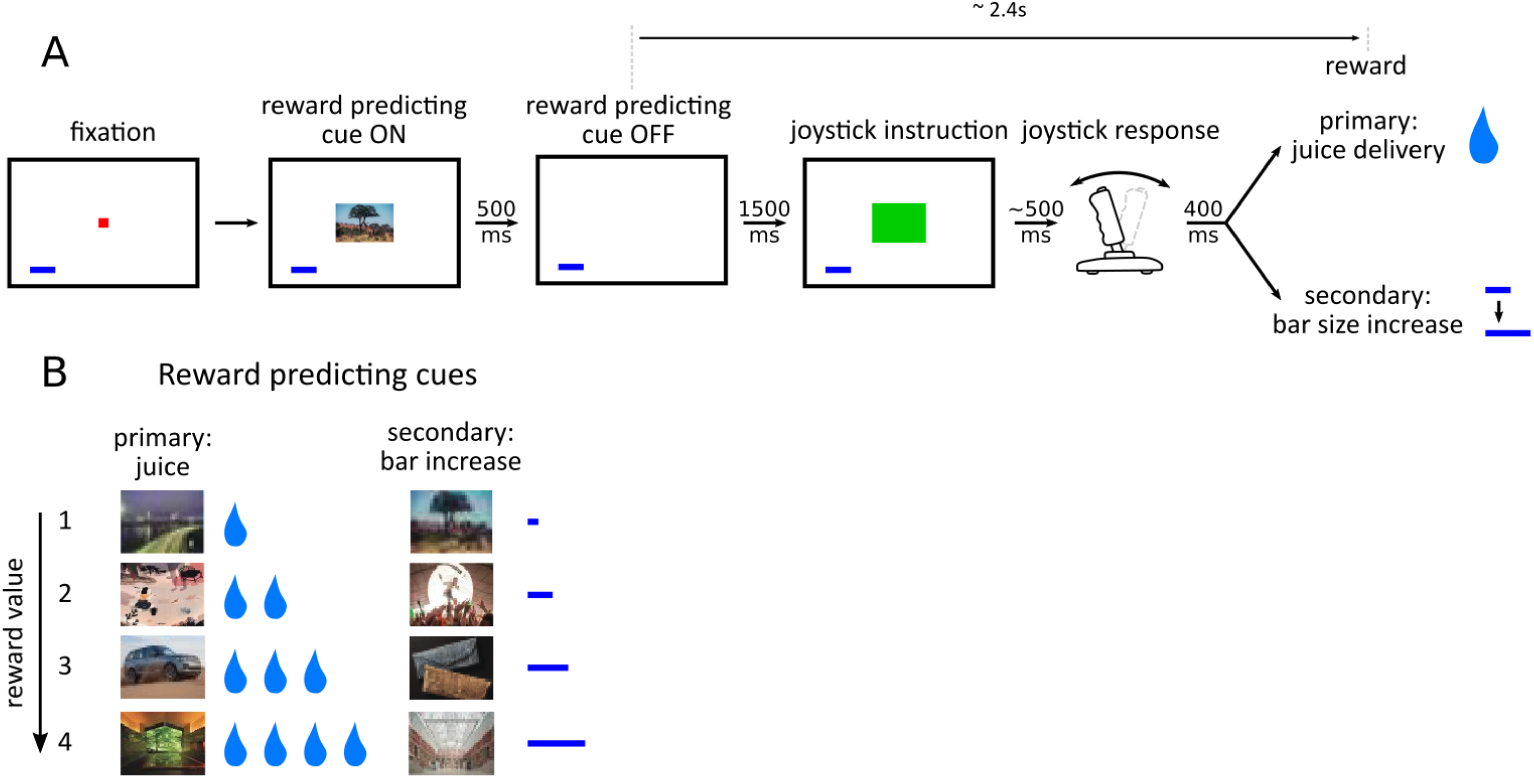
Value-based decision making task. **A** - Monkeys initiated a trial by fixating a point in the center of the screen. A reward predicting cue appeared that the subject was required to fixate for 450 ms. After a 1500 ms delay, one of two possible images instructed the monkey to move a joystick right or left. Contingent on a correct joystick response, monkeys received a reward either in the form of juice (primary) or an increase of a reward bar (secondary), which was constantly displayed on screen (blue bar in figure). Note that the presentation of the reward predicting cue and the delivery of the reward are separated in time by more than 2 s. **B** - A total of 8 reward predicting pictures covered the combinations of 4 possible values, and 2 reward types.

Monkeys were presented with choice trials (∼20% of trials), in which the animals could choose between two stimuli presented simultaneously on the screen. Choices were made by fixating the stimulus of their choice, and the associated reward would be delivered at the end of the trial. Both monkeys learned to perform the task optimally by selecting the stimulus associated with higher values [Rich and Wallis2016]. We focused on the remaining trials that included a single cue only. On these single picture trials, reaction times in the unrelated joystick task decreased as the value of the stimulus increased, providing evidence that monkeys maintained an expectation of the upcoming reward across the intervening time interval [Rich and Wallis2016]. Here, we investigated the representation of value during this period between the onset of the reward predicting cue and reward delivery, which lasted ∼3s, and included a ∼2.4s delay after the cue offset.

Single and multi units were recorded in the orbitofrontal (OFC) and anterior cingulate cortex (ACC) of the two subjects while they performed the decision making task across 24 (monkey M) and 20 (monkey N) sessions (Table 1).

**Table 1:** Number of recorded units with an average activity per session above 1Hz.

| Monkey | Single |  | Multi |  | Total |
| --- | --- | --- | --- | --- | --- |
|  | M | N | M | N |  |
| OFC | 259 | 192 | 170 | 177 | 798 |
| ACC | 114 | 72 | 81 | 48 | 315 |

Multi and single units from all recording sessions and both monkeys were pooled together to create 2 populations of 798 and 315 neurons recorded from the OFC and ACC respectively. Basic unit and decoding results did not differ across monkeys (Sup. Fig. 1) so data were pooled.

### Encoding and decoding of value in unit activity

To assess the encoding of task variables by the neurons in these two regions, we regressed the firing rate of single and multi units on variables value, type and their interaction value x type and tested the contribution of these variables with an ANOVA. We considered value as a categorical variable, as some units activated in a non-linear fashion, for example by responding only to a specific value and could not be modeled with a simple linear regression (Sup. Fig. 2).

**Figure 2:**
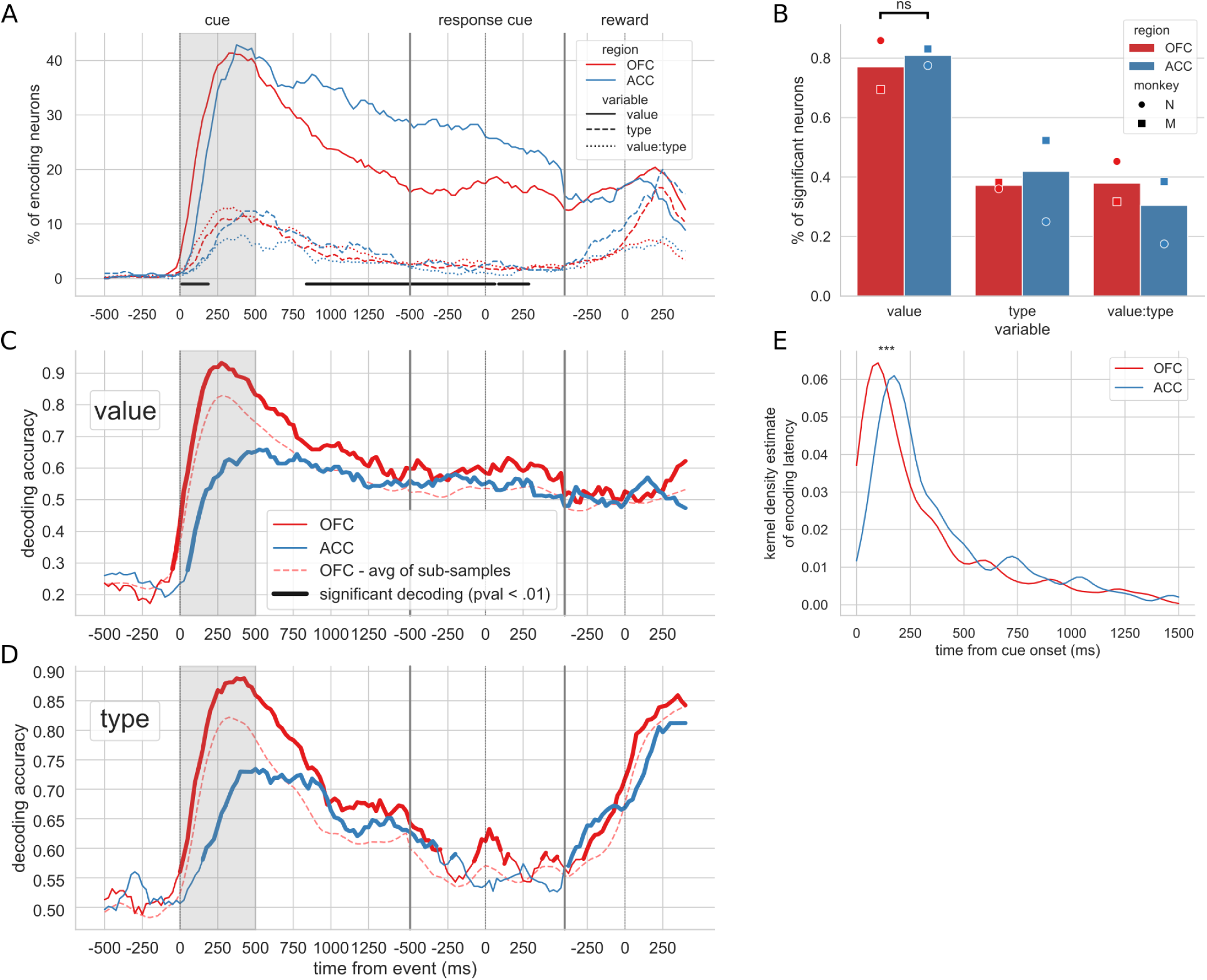
Encoding and decoding of value. **A** - Percentage of units encoding the different variables at each point in time in both OFC and ACC. Significant difference in value encoding neurons with Chi^2^test is indicated by thick black lines at the bottom (p<0.01). **B**. - Percentage of units encoding the task variables at any point in time from the onset of cue presentation to the delivery of the reward. **C** and **D** - Decoding accuracy of value and type, respectively, with a ridge regression classifier that was trained and tested independently on each time point. Significant decoding is shown by the thick portions of line and corresponds to a p-value < .01 with a 2000-permutation test. Dashed red line is the average of 200 sub-samples of OFC data with the same number of units as in ACC for comparison. **E** - Kernel density estimate of unit value encoding latencies. OFC latencies are significantly shorter (see main text).

Similar to previous reports [Rich and Wallis2017], the vast majority of OFC and ACC units encoded value at some point after the reward predicting cue and before the reward (77.7% for OFC and 80.2% for ACC, Fig. 2B). The proportion of neurons encoding value peaked after the presentation of the reward predicting cue and steadily decreased until the reward (Fig. 2A). The type of reward was only weakly encoded compared to value, with only half the proportion of units encoding it at any time (37.1% for OFC and 38.7% for ACC, Fig. 2B). No difference between ACC and OFC was found in the proportion of units encoding value at any point during the epoch of interest (Fig. 2B, Chi^2^test of independence, p=0.18). The proportion of units encoding value was higher in ACC during the delay (Fig. 2A), however this difference is mostly driven by the data from one monkey and is not significant in the other subject (Fig. Sup. 1A). In accordance with previous findings [Kennerley and Wallis2009], the relative onset of value encoding was earlier in the OFC compared to ACC (Fig. 2E). Latencies in the OFC were significantly shorter (Kruskal-Wallis on latencies [stat=19.4, p=1.05*10^−5^] and permutation tests confirmed mean-latency difference [mean=72.6ms, p=2.9*10^−3^]). This result was replicated in each monkey’s data independently (Fig. Sup. 1C).

Population decoding of value with a ridge classifier elicited a peak in accuracy similar to the peak in the proportion of encoding units after the presentation of the delay followed by a stable significant decoding until reward (Fig. 2C, significance threshold set to p<0.01 with 2000 permutations, replicated in each monkey, Sup. Fig. 1B). A similar result was observed for decoding type (Fig. 2D), although decoding accuracy rapidly dropped and reached non significance around the joystick task. The first significant bin decoding value and type was 100ms and 150ms earlier respectively in OFC compared to ACC and peak decoding was earlier and higher in OFC compared to ACC (for value/type: 93.2%/88.8% in OFC at 275ms/425ms, 65.9%/73.4% in ACC at 525ms/500ms). To fairly compare decoding accuracy between the two regions, we randomly sub-sampled the OFC population to match the number of ACC units 200 times. The average accuracy of the sub-sampled populations was still higher in OFC than ACC, indicating that the higher number of units in OFC alone cannot explain the difference in accuracy. Together, these results show that value is strongly encoded in these prefrontal regions and can be significantly decoded throughout the delay. Because of the lower incidence of reward type encoding and low type decoding accuracy, the rest of the study focused on the representation of value.

### Persistent versus sequential encoding

The total number of neurons encoding value during the delay was much higher than the number encoding at any particular point in time, indicating that value was represented by different neurons throughout the delay. Given this, we next sought to understand the contribution of individual units to the population dynamics. A common method for doing this is sorting neuron responses. For example, to demonstrate that a neural population uses a sequential activation scheme, sorting the neurons by their peak activation typically displays a tiling of the delay. To ensure that sequential encoding of value is a robust feature of OFC and ACC activity, with split the data in half into training and testing trials. We sorted the units based on the time of peak value encoding during the delay with the training half and found that both OFC and ACC value coding tiled the delay, such that different subsets of neurons were most selective at different times (Fig. 3A). We then sorted the testing data with the peak encoding bin of the training data to assess its consistency (Fig. 3B) and found that the tiling of the delay is present but less clear in the testing data. Indeed, the peak encoding bin of the training and testing data set was significantly correlated, which is an expected observation with sequential encoding, though many neurons were far from the diagonal, hinting at a non-uniform distribution of peak encoding times across the delay. Such results could be used to support the hypothesis of a sequential representation of value in both areas, consistent with conclusions drawn in other studies [Pastalkova et al.2008], although the mere firing rate of units is often used (see Sup. Fig. 3A and D for firing rates). However, sorting the neurons by the proportion of the delay during which they encode value offers a different picture. Fig. 4A and B show that while some units have time-limited encoding of value, another portion continuously encode value throughout the delay supporting the notion that there is a stable representation of value in the population. A congruent result was obtained by clustering the pattern of average firing rate of each neurons in two clusters (Sup. Fig. 3B and E), such that one cluster showed a pattern of increased firing rate for the duration of the delay. This approach gives the impression that a large proportion of units display persistent activity. However, clustering the average firing rate of units with 6 clusters uncovered more diverse profiles of activity that nonetheless follow basic principles: phasic activity followed task events with stable or monotonically evolving activity between events. Thus, depending on the sorting method employed, the dynamic and sequential or stable and persistent hypothesis can be defended. Overall, most units covered a relatively small portion of the delay. A reverse cumulative distribution of the number of units encoding value showed a sharp decrease in the fraction of the delay covered (Fig. 4B). Only 6.9% and 10.2% of units encoded value for more than half the delay in OFC and ACC respectively. Further, 80% coverage of the delay was found in roughly 1% of the units (OFC: 0.75%, ACC: 1.59%). The distribution of delay coverage by encoding units did not differ between OFC and ACC (Kolmogorov-Smirnov test, KS=0.079 and p=0.90).

**Figure 3:**
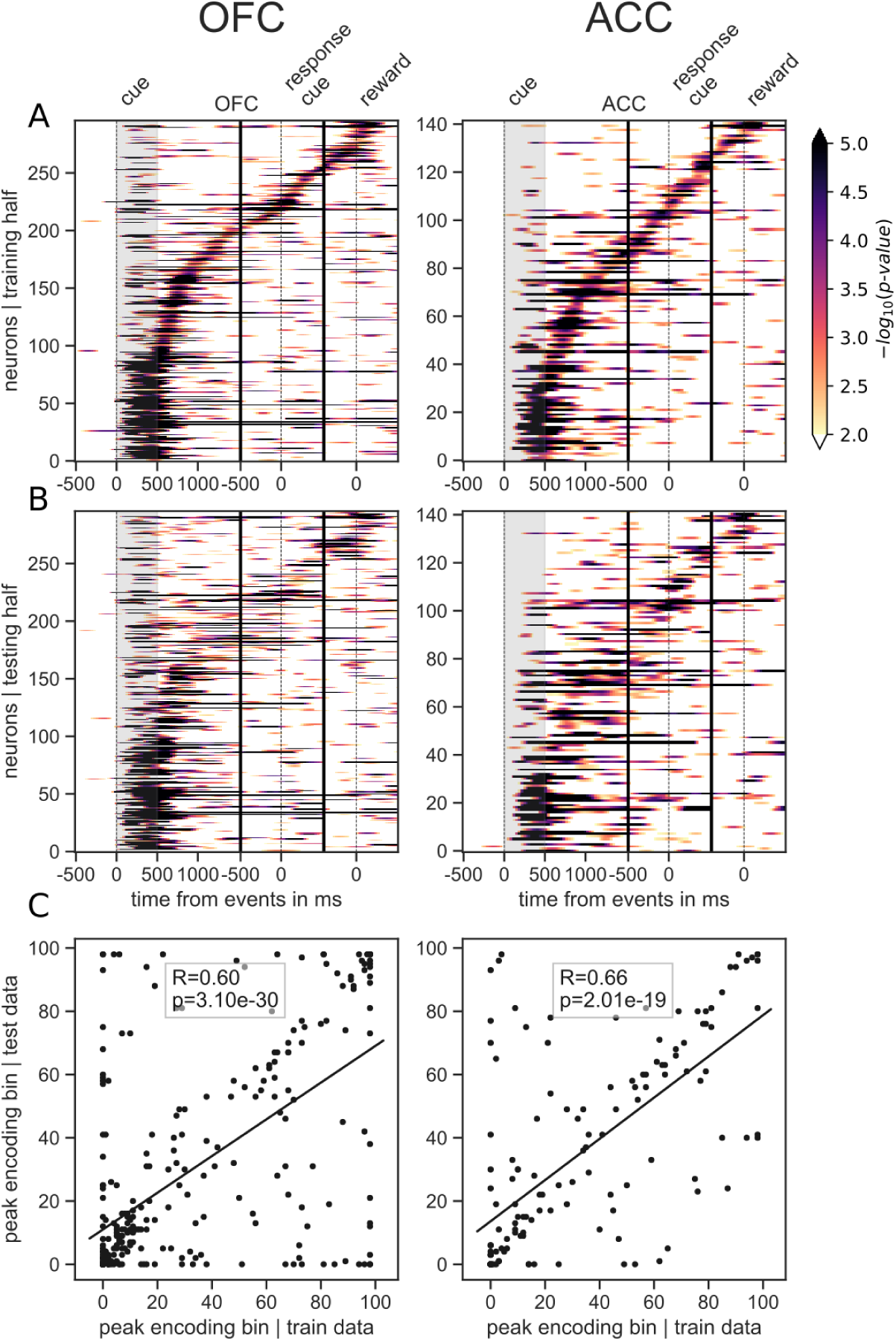
Tiling of the delay by value encoding units. **A** - Tiling of individual units according to their peak encoding of value across the delay in both OFC and ACC with the training half of the trials. Colors represent the negative log of the encoding p-value (ANOVA, F test on value). The measure is bounded between 2 and 5 for visualization and corresponds to p-values ranging from 0.01 to 10^−5^. **B** - Tiling with the testing half of the trials, with units sorted according to the training half of the trials, to show consistency in sequential encoding. Note that only units that had significant encoding in both the training and testing dataset were included, hence the lower number of units. **C** - Scatter plot of the peak encoding bin for training versus testing half of the data. Graphs show the Spearman correlation coefficient R with associated p-value p and trendline.

**Figure 4:**
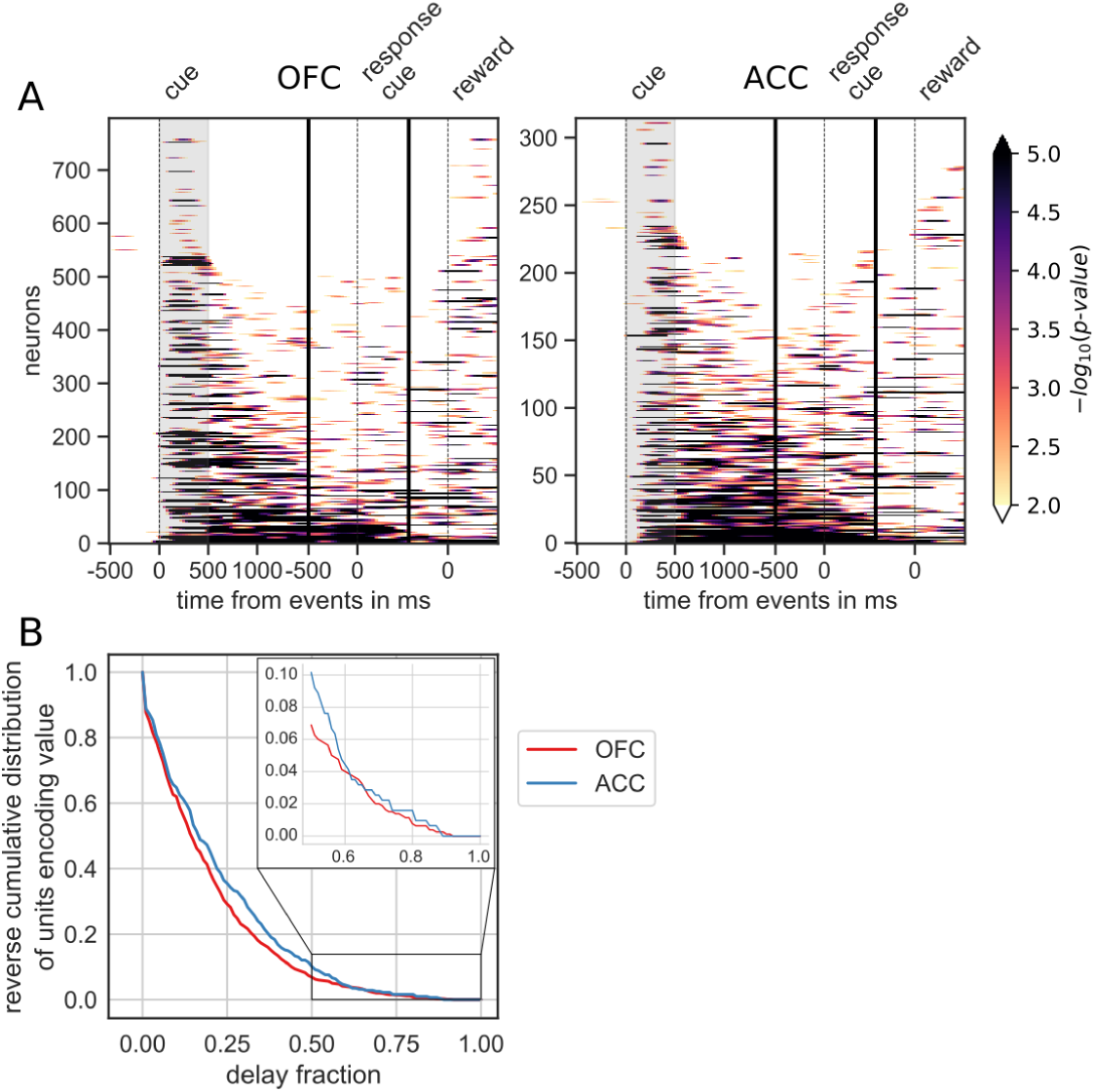
Spanning of the delay by value encoding units. **A** - Units sorted by the duration of encoding of value. Colors represent the negative log of the encoding p-value (ANOVA, F test on value). The measure is bounded between 2 and 5 for visualization and corresponds to p-values ranging from 0.01 to 10^−5^. **B** - Reverse cumulative distribution of units encoding value as a function of the fraction of the delay covered. This graph shows the proportion of units encoding value for at least the fraction of the delay indicated on the x-axis (no significant difference between regions, see main text).

### Population Dynamics

Decoding analyses allow us to assess how well value information can be extracted from the population as a whole, but standard approaches that find unique solutions to optimally decode value at each point in time say little about the dynamics of a representation. In addition, our results show that value encoding among units can paint contrasting pictures at the level of population representation, i.e. as both dynamic and sequential or stable and persistent. Therefore, to explore representational dynamics across time, we used cross-temporal decoding (CTD), in which a decoder is trained and tested at every possible pair of time bins in the delay [Stokes et al.2013, Astrand et al.2014, Stoll et al.2016, Meyers et al.2008]. In this context, the classification accuracy is a proxy for the similarity in value representation between any two time points. We can then produce an accuracy matrix consisting of each training/testing time bin pairs. When the training and testing time are identical, we obtain a classical decoding procedure as shown in figure 2C, which corresponds to the diagonal in the CTD accuracy matrix.

Using this method on the OFC and ACC populations, value representation dynamics are not clearly dynamic or stable (Fig. 5A). OFC evidenced a somewhat stable representation from 500 ms after the reward predicting cue (around cue offset) until the response cue, but the representation remained dynamic before and after this period as the accuracy was mostly confined to the diagonal. ACC accuracy was lower overall but showed similar patterns. While significant decoding appeared to spread from reward cue to after the response cue, accuracy was not homogeneous and remained strongest along the diagonal, indicating a rather dynamic representation of value.

**Figure 5:**
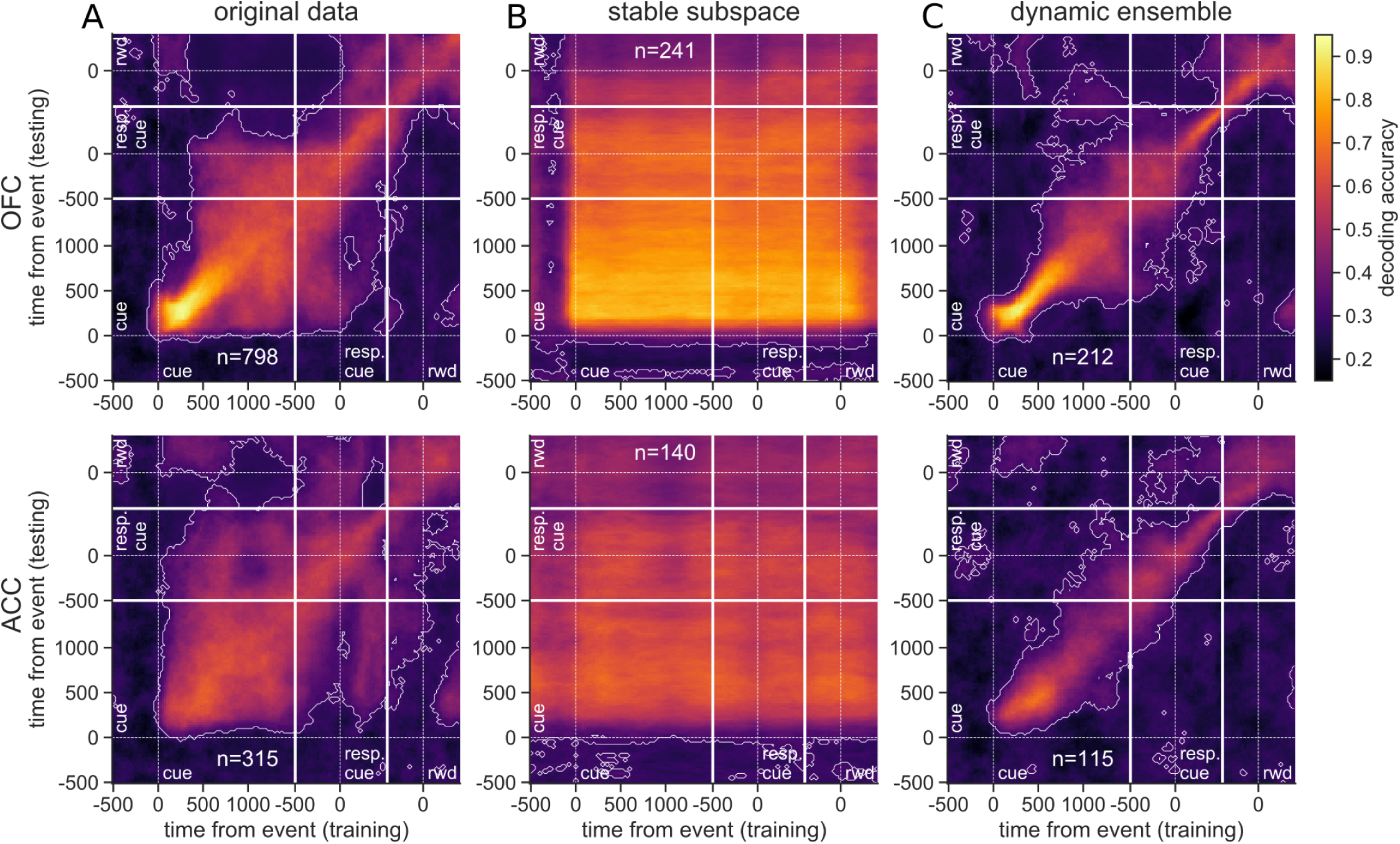
Accuracy of cross-temporal decoding (CTD) of value with different methods. Training and testing of a ridge classifier at different time bins across the delay. CTD was applied to (**A**) original data, (**B**) a stable subspace obtained from the combination of a value subspace and an ensemble method where units are iteratively removed to obtain maximum accuracy, and (**C**) on a dynamic ensemble where units were iteratively removed from the ensemble to maximize a dynamic score. Thick white lines indicate junctions between the three successive epochs (cue presentation, joystick task and reward delivery). The thin dashed white lines indicate the reference event of each epoch (reward predicting cue onset, response instruction, reward). Contour curves indicate areas with a p-value lower than 0.01 with a 2000 fold permutation test. (rwd = reward, resp. cue = response cue)

This dynamic pattern was also reflected in the population trajectory speed obtained by calculating the distance between two successive time bins in the neural space (Fig. 6). The reward predicting cue triggered an increase in speed that was followed by a period of low speed, indicative of relatively stable population activity, until the joystick instruction cue triggered another period of rapidly changing activity. Thus strongly dynamic bouts of activity occur with changes in task stimuli, and in the absence of such changes, delay activity becomes quite stable.

**Figure 6:**
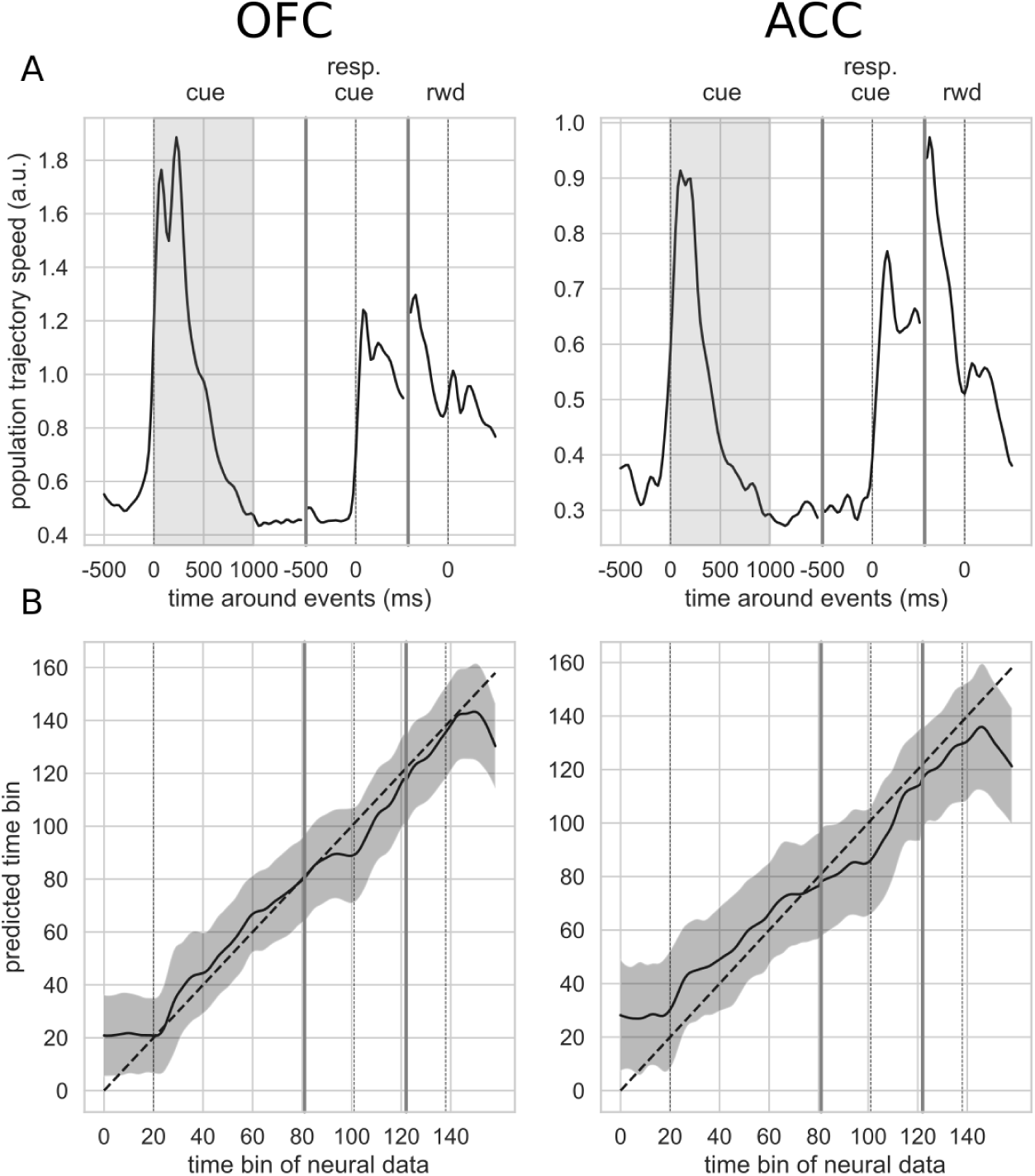
Trajectory speed and regression of time. **A** - Speed (rate of change) of the average activity trajectory in the neural space defined by the full population of neurons, in OFC (left) and ACC (right). Note that the difference in scale between OFC and ACC (y-axis) is due to the difference in the number of units in each population. **B** - Regression of the time bin from population activity with a simple linear regression. The black line and shaded area represent the average and standard deviation of leave-one-out cross-validation results. The dashed line corresponds to ground truth. For more temporally accurate results, the firing rate of each unit was estimated with a 50 ms standard deviation Gaussian kernel instead of the 100 ms used in the other analyses.

### Stable representation of value

The basic CTD method relies on a decoder that weighs on the most discriminating features (i.e. neurons) at a particular point in time. If a sub-population strongly encodes value at one time, the decoder will attribute stronger weights to these units. However, if this sub-population changes representations or ceases to encode value, the decoder accuracy will drop. Our results suggest such temporally local representations that change across time, but this could simply be due to inhomogeneous representation of value at the population level. That is, the strongest representation at one time point may not generalize well to other time points. This would not exclude the possibility that a stable representation of value exists within the population, because the classifier is not optimized to find a decoding plane that is consistent throughout the delay.

To determine whether a linear combination of neural activities can be used to extract a stable representation of value, we combined two additional methods to optimize CTD. The first is a subspace method inspired by [Murray et al.2017], in which data are projected into a subspace that maximizes value representation while lowering the influence of temporal dynamics (see Methods). This was applied to data from the onset of cue presentation to the delivery of reward. A three-dimensional subspace was obtained, and data projected onto this subspace was used to decode value, using CTD in the same manner as was done for the full population. Our cross-validation approach ensured that test data were not used to generate the subspace, as it would introduce a bias in decoding results (see Methods).

Repeating the CTD procedure with data projected onto the subspace demonstrates that stable value representations can be extracted from the same population data (Sup. Fig. 4A). Though significant decoding using CTD did not fully span the cue to reward period in OFC, higher accuracies were more widespread. ACC accuracy, which also spanned a larger portion of the delay, covered nearly all of the epoch between cue and reward using the subspace approach.

Although a higher unit count is expected to produce higher decoding performances, in the case of CTD better performances might be obtained by pre-selecting the neurons that most participate in the stable decoding over time. For this purpose we derived an ensemble method inspired by [Backen et al.2018] that iteratively selects units based on their contribution to decoding accuracy. Essentially, we started by removing each unit independently to find the n-1 ensemble that most increased the overall CTD accuracy, then removed a second unit to find the best n-2 ensemble and so on until only one unit was left, and then selected the ensemble that maximized decoding accuracy. This procedure markedly improved stable decoding across time, with significant decoding spanning all time pairs after the reward predicting cues (Supplemental Fig. 4C). The ensembles that best optimized the CTD accuracy in both regions contained only a fraction of the units from the full population (212 out of 798 units for OFC and 115 out of 315 units for ACC). Greater numbers of units did not improve performance, likely because many of these modified their value encoding over time and did not provide a stable signal.

While the subspace and best ensemble approaches both found somewhat stable value representations, the most stability with highest accuracies was obtained with a combination of these methods. The subspace procedure was applied to each ensemble selected by the iterative ensemble method. The result is a strikingly stable CTD in both regions, albeit with slightly lower accuracy in ACC compared to OFC (Fig. 5B). This demonstrates that when CTD is applied to a neural population, the representation may appear to be partially dynamic, yet it is possible to find a subspace of the population activity that defines a completely stable representation across time. In addition, combining subspace and ensemble procedures is a promising method to extract extremely stable representations of task variables. These results were replicated with data from each monkey independently (Sup. Fig. 5).

### Dynamic representation of value

Since we found that stable value representations could be extracted with appropriate methods, it is logical that dynamic representations can also be identified. To do this, we first defined a “locality” measure to quantify how local a representation is based on the decoding accuracy over time. We fit a Gaussian curve to the decoding accuracy obtained from training at a given time bin and testing on all time bins. The height of the Gaussian divided by its standard deviation defined a score of temporal locality. The measure was defined as the average of the scores computed from the accuracy curves obtained across all training time bins. A dynamic representation of value was obtained by selecting the neurons with the ensemble method described above that maximized this measure. The resulting CTD accuracy displays the main feature of a strongly local and dynamic representation of value, where higher accuracy is confined to the diagonal (Fig. 5C). With the original full population, value representation in the OFC was rather stable in the middle of the delay, but the dynamic ensemble shrunk the higher accuracy block to constrain it to the diagonal. The most striking result was obtained from the ACC population, where significant accuracy that extended from the reward-predicting cue presentation to the response instruction with the full population was well confined to the diagonal with the dynamic ensemble, leaving only a very local representations. The difference in the OFC population might be explained by slower population activity dynamics during the delay, after the transient phasic activity elicited by the reward-predicting cues. Regardless, this result demonstrates that a dynamic subspace encoding local value representations can be extracted from the same neural population that encodes value in a very stable way. Together, these results support the idea that a basic CTD applied to a neural population yields a limited view of mixed neural dynamics.

Since the dynamic and stable ensembles yielded opposite dynamics, we expected limited overlap between them. Indeed, among the 798/315 units in OFC/ACC populations, there were 139/71 units in the stable ensemble and 212/115 units in the dynamic ensemble but only 22/17 units were part of both ensembles, which is less than expected by chance (Chi^2^test of independence, p=0.002/p=0.018).

To further demonstrate that the activity in both these areas is dynamic and supports the encoding of time, we regressed the index of time bins from 500 ms before the reward predicting cue onset until 500 ms after reward delivery with a simple linear regression from the neural population activity (Fig. 6B). Indeed, time can be predicted accurately, supporting the notion of a robust representation of time in the neural population activity.

### Relationship between value encoding and decoding dynamics

It is logical to expect that units that strongly encode value over long periods of time should participate more in the stable subspace. To test this, we correlated different encoding measures of units with their contribution to the value subspace and stable ensemble (Spearman correlation based on rank, see Methods). We defined the encoding strength score as the average of the negative log p-value of each unit across the delay, and then found the fraction of the delay when the unit significantly encoded value (p-value < 0.001, for at least 7 bins in a row). First we correlated each of these measures with the unsigned value subspace weights, since a stronger weight reflected a higher influence on the subspace irrespective of its sign. Both measures were highly correlated with subspace weights (Fig. 7A, left and middle) indicating a clear link between encoding strength and duration and contributions to the value subspace. However, units might also change their pattern of value encoding during the delay. To quantify this, we defined an additional stability measure that includes the duration, strength and stability of value representation by penalizing cases where the difference in average firing between a pair of values changes sign (see Methods for details). This measure was also highly correlated with the value subspace weights (Fig. 7A, right graph). Overall, these measures describe the qualities of activity that promote selection to the value subspace. Indeed, the subspace method relies on trial and time averages of unit activities, so units having strong and long value encoding are more likely to be selected by this averaging method.

**Figure 7:**
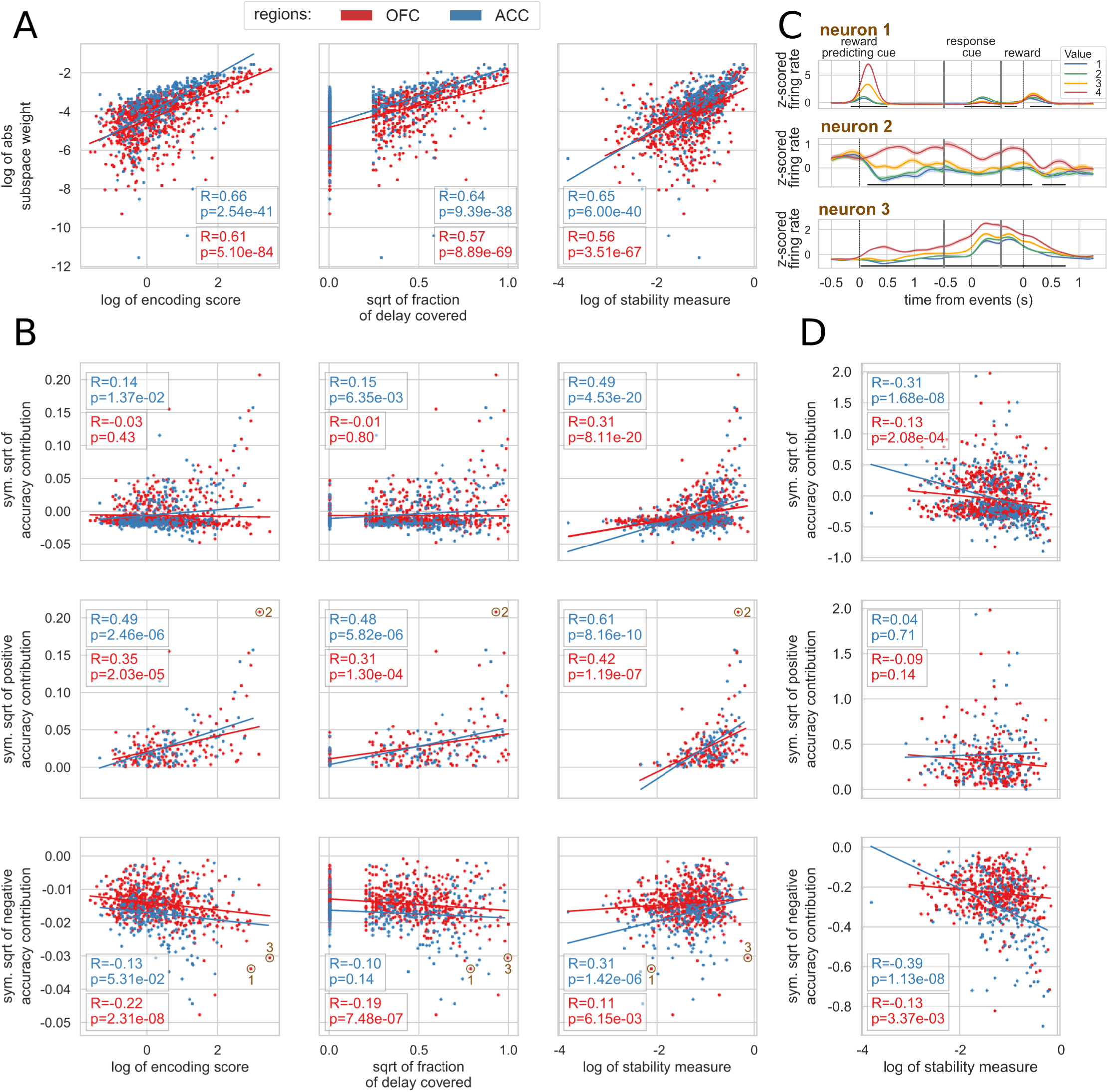
Correlation between encoding and decoding measures. **A** - Correlations between encoding measures and subspace weights. **B** - Correlations between encoding measures and CTD accuracy contribution. Top row includes all contributions, middle row only positive contributions and bottom row only negative contributions. **C** - Z-scored firing rate of two example neurons negatively contributing to the stable ensemble (1 and 3) and one neuron contributing positively (2). Individual data points corresponding to these neurons are circled in **B**. **D** - Correlations between stability measure and locality measure contribution. All figures show Spearman correlations. sqrt=square root; negative contributions were transformed with a symmetrical function around the origin based on square root: 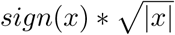 Transformations applied to the data had no effect onhe correlations which are rank based (see Methods).

While strength and duration of encoding predicted which neurons contributed to the stable subspace, they poorly predicted which neurons were selected by the ensemble method. To explore the contribution of each unit to the ensemble, we defined an accuracy contribution score based on the change in mean accuracy across all time bin pairs in the CTD elicited by removing a unit from the ensemble. This accuracy contribution score for each unit was either not or weakly correlated with encoding strength and duration among OFC units, and even less among ACC (Fig. 7B, top row). However, accuracy contribution was correlated with the stability score in both regions. Because neurons with positive and negative contributions created asymmetrical distributions, correlations were repeated on each of these independently. Interestingly, in both regions, encoding strength and duration were correlated with positive contributions to the stable ensemble (Fig. 7B, middle row), however only OFC was significantly anti-correlated with negative contributions (Fig. 7B, bottom row). Positive contributions were even more correlated with the stability measure, and contrary to the other encoding measures, positively correlated with negative contributions. Therefore, neither the encoding strength nor duration measures by themselves fully describe how much a unit contributes to stable ensemble decoding. A strongly encoding unit might encode value for a limited time and/or reverse its encoding (e.g. Fig. 7C neuron 1), and a unit covering a large part of the delay might also change over time and not provide a stable representation. Conversely, units with stable encoding contribute more to the ensemble (Fig. 7C neuron 2). However, since decoding relies on the interaction between many neurons, these simple encoding measures of individual units cannot fully explain all the ensemble contributions. For example, neuron 3 in 7C strongly encodes value for the whole delay, but contributes negatively to the stable ensemble.

Finally, contributions to the dynamic subspaces were negatively correlated with stability, as expected (Fig. 7D) among both positive and negative contributions, though the anti-correlation appeared to be mostly driven by negative contributions. Thus, neurons that represent value in a stable way for long periods of time are unlikely to be contributing to a dynamic subspace designed to extract temporally limited value representations.

## Discussion

In this study, we shed light on the complex dynamics of value representations in prefrontal cortex. While value representations have been found in the activity of ACC and OFC e.g. [Rushworth and Behrens2008], this is, to our knowledge, the first study of their dynamics during a short delay. We show that targeted methods elicit seemingly opposite results that do not lend themselves to straightforward interpretations with respect to the current theoretical frameworks common in the WM literature. Unit encoding shows features of both persistent and sequential activity, depending on which feature the analysis method is designed to extract. Similarly, targeted methods can extract either a stable subspace or a temporally local representation at the population level. These results, along with recent studies of the dynamic nature of delay activity in the prefrontal cortex [Spaak et al.2017, Murray et al.2017, Meyers2018], argue in favor of a more nuanced view than the pure persistent/stable, activity silent/dynamic or sequential activity hypotheses.

Here we propose that mixed dynamical regimes, with both time and temporally insensitive information concurrently encoded in the same neural population, can provide a richer substrate to face varying task demands. On one hand, the main function of WM-related processes is to temporarily maintain information not available in the sensory environment so it can be manipulated or retrieved later. A stable neural representation, as modeled in networks with strong attractor dynamics, is an efficient means of holding such information online. Stability is particularly useful when information must be extracted at an unspecified time. In this case, a target population could use the same method to reliably read out an unchanging representation whenever it is required. Conversely, representing relevant information in a time sensitive manner is also critical, since most behaviors are organized in time, whether they are internally generated or aligned to external events [Meyers2018, Cueva et al.2019]. For example, our task includes a recurring sequence of events (fixation point, reward predicting cue, delay, joystick instruction cue, etc…), which the subjects have learned. By representing time, they can anticipate the next event and prepare to either process a task relevant stimulus or produce a motor output. Such preparatory processes can help optimize task performance and maximize the amount of reward the subject can receive. Given this perspective, a key question that remains is whether dynamic value representations are merely the result of mixing time and value information that are independently required to perform the task, or whether specific combinations of these are necessary to perform the task optimally. Future work probing the roles of these features in artificial neural networks could address this question.

The debate about persistent versus dynamic representation in WM tasks has relied heavily on modeling experiments. Neural network models of persistent activity have historically been implemented as fixed point attractors when the remembered information comes from a discrete number of items [Amit1995, Compte et al.2000] or bump/line attractors in the case of parametric working memory [Machens et al.2005, Seung1996]. Another neural network framework, with random recurrent connections referred to as reservoir computing, subsequently demonstrated that short term memory is readily obtained from generic networks in which different inputs (stimuli) elicit distinct trajectories thereby encoding stimuli for a short duration with highly dynamic signatures [Maass et al.2002, Jaeger2001]. While these dynamic properties appear irreconcilable, they are not necessarily exclusive and can be combined in a single network, as demonstrated by more recent models [Maass et al.2007, Pascanu and Jaeger2011, Enel et al.2016]. Such networks can provide mixed dynamics similar to those observed in the present task, suggesting that they are better models for short-term value maintenance in OFC and ACC.

Network modeling experiments also provide theoretical explanations for the degree of persistent versus dynamical activity. Specifically, some emerging theories suggest that this trade-off is a function of task requirements instead of the network architecture. For example, [Masse et al.2019] argue that tasks requiring manipulation of information held in memory recruit more persistent activity. [Orhan and Ma2019] explored several task parameters, among which fixed delay duration and higher temporal complexity elicited more dynamical activity. However, it should be noted that very little is known about how the choice of network training algorithm affects the dynamical regime of these networks. Similarly, little is known about the effect of training on neural representations in experimental subjects such as monkeys. Indeed, it is likely that the degree of training a subject receives in a cognitive task will influence the neural representations recorded in that task. For instance, the degree of stable versus dynamic representation might vary depending on whether the subject is over-trained on the task or on particular stimuli in the task, such that over training might shift weak and dynamical activities toward more stable and persistent ones. In this context, the work of [Barak et al.2013] demonstrate how different model architectures and training methods could account for the gradual morphing of population activity from transient to attracting dynamics and could explain the variety of dynamics observed in recorded prefrontal activity.

Among the dynamic representations of value, we found some evidence for sequential organization within our neural populations. Sequential activity is the successive, short-duration activation of individual neurons that are thought to either pass information to bridge a delay or represent the passage of time in the absence of external stimuli [Pastalkova et al.2008, Rajan et al.2015]. Such activity is not necessarily synonymous with dynamical activity, as the former is a specific and extreme case of the latter. Beyond the mere representation of time, sequential unit activity has been associated with a variety of functions, including the temporal segmentation of memories in the hippocampus [MacDonald et al.2011, Eichenbaum2014] and WM in parietal cortex and the rat medial PFC [Harvey et al.2012, Fujisawa et al.2008]. To the best of our knowledge, this type of sequential activity has not been reported in prefrontal regions of primates performing WM tasks, potentially because it has not been the focus of analyses to date. Consistent with the heterogeneous nature of our neural population, only some neurons showed evidence of reliable sequencing, while other neurons’ activation and encoding was not short and uniform. Sequencing appeared most prominent early in the delay, and may coincide with the brief, highly dynamical epoch immediately after the presentation of a stimulus to be remembered that has been found in this and other studies. This dynamic period is typically followed by a more stable representation until the end of the delay [Stokes et al.2013, Murray et al.2017, Barak et al.2010, Wasmuht et al.2018, Cavanagh et al.2018]. It has been proposed that the dynamical epoch following the memorandum presentation is the result of stimulus processing, which in some tasks corresponds to the transformation of the stimulus into task relevant information [Stokes et al.2013], and our results suggest this process may involve more reliable sequencing among neurons. More generally, it appears that task relevant inputs to a brain region will trigger local processing that is reflected in transiently increased and dynamic activity as was demonstrated by a reservoir model mimicking ACC activity dynamics [Enel et al.2016].

One challenge for attractor networks modeling WM tasks has been maintaining stable representations in the presence of distractors or intervening task demands. In the current study, monkeys had to perform a joystick sub-task before receiving reward, and CTD with the original neural populations yielded stable value representations until the presentation of the joystick instruction cue. Similar dynamics have been observed when a distractor perturbs the representation of a spatial cue held in WM [Parthasarathy et al.2017, Cavanagh et al.2018]. The initially stable representation might be the result of unperturbed network activity that persists until a task relevant input (the response instruction cue) impacts the networks’ dynamics. However, here we show that analyses designed to identify stable representations can, indeed, extract a subspace of the same population activity that was stable across the intervening joystick task. Interestingly, value encoding and basic decoding remained largely unaffected during the intervening task, unlike results previously reported in a dual task experiment. Here, when monkeys solved attention and memory tasks concurrently [Watanabe and Funahashi2014], selectivity for components of either task diminished at the level of single neurons and population activity. Similarly, intervening stimuli or distractors presented during the delay of a WM task disrupt the representation of memoranda [Lebedev et al.2004, Warden and Miller2007, Jacob and Nieder2014]. The discrepancy between these and our results might lie in the non-overlapping nature of value information and joystick task. In our case, the task and stimuli were independent, whereas in these other studies, a stimulus of the same nature as the memoranda (e.g. a visual stimulus) was presented, perhaps triggering a conflicting representation.

Finally, a notable conclusion from this study is that OFC and ACC have similar dynamics that represent value throughout a delay. Despite this overall similarity, a few key differences were found. First, value was encoded less than 100ms earlier in OFC compared to ACC. Second, value was more easily decoded in OFC immediately after the reward predicting cue, while a higher proportion of ACC neurons encoded value during the delay. These results argue in favor of the notion that value information is represented and processed earlier in the OFC than in ACC [Kennerley and Wallis2009]. Nonetheless, the mixing of dynamic regimes appears quite similar between these two areas, so that both encode time sensitive and insensitive representations of value. Therefore, the underlying neural mechanisms responsible for encoding value over time are likely similar in OFC and ACC.

Overall, using analysis methods designed to extract different signal dynamics, we found that all co-exist to some extent within the same neuronal populations that represent the same task-related variable. Rich representations of expected values are critical for adaptive learning, and the mixed dynamics reported here could play an important role in conveying both general and temporally specific value expectations to areas involved in optimizing goal-directed behavior.

## Methods

### Behavior and neurophysiology

The behavioral task has been previously described in [Rich and Wallis2016, Rich and Wallis2017]. Two head-fixed male Rhesus macaque monkeys (Macaca mulatta), aged 7 and 9 years, weighing 14 and 9 kg at the time of recording, were trained to perform a value-based decision making task in which they chose between visual stimuli associated with rewards. In the present study we focused on single cue trials only. All procedures were in accord with the National Institute of Health guidelines and recommendations of the University of California at Berkeley Animal Care and Use Committee. Subjects sat in a primate chair, viewed a computer screen and manipulated a bidirectional joystick. Task presentation and reward contingencies were controlled using MonkeyLogic software [Asaad and Eskandar2008], and eye movements were tracked with an infra-red camera (ISCAN, Woburn, MA).

A total of 8 pictures (∼2° × 3° of visual angle) comprised the set of stimuli associated with rewards. Pictures were selected randomly from this set on each trial. Four pictures predicted the delivery of juice reward (0.05, 0.10, 0.18, 0.30 ml), and four predicted that the length of a reward bar always present on the screen would increase by a set increment. Subjects were previously trained to associate the length of the bar to a proportional amount of juice obtained every 4 trials. Associations between cue and reward were probabilistic, so that on 4 out of 7 trials the type and amount of reward was consistent with the cue. On 1 out of 7 trials, the type of reward was different, on 1 out of 7 trials, the amount was randomly picked to be different, and on 1 out of 7 trials, both the amount and type differed from what the cue predicted.

Single cue trials were randomly interleaved with choice trials. Because the different reward values for primary (juice) and secondary (bar) outcomes were titrated, monkeys almost always chose the target associated with a higher value irrespective of the type of reward [Rich and Wallis2016].

During the delay between the reward predicting cue and reward delivery, monkeys were required to move a joystick either left or right depending on a visual cue, which we refer to as response instruction to avoid confusion with the reward predicting cue. Reward delivery was contingent on a correct joystick answer, and reaction times were inversely correlated with the size of the expected reward [Rich and Wallis2016].

Electrodes were lowered at the beginning of each session in the OFC (areas 11 and 13) and dorsal bank of the ACC sulcus (area 24). Recorded units were not screened for selectivity, but those with average activity lower than 1Hz across the session were excluded. Further details on behavior and recording methods can be found in previous publications [Rich and Wallis2016, Rich and Wallis2017].

Firing rates of each unit were estimated with a 100ms-SD Gaussian kernel convolution over the spike train of entire recording sessions. The resulting firing rate was divided into epochs around the reward cue, the response (joystick) instruction cue and reward delivery, then binned into 100ms bins every 25ms, so that there were 40 bins for 1 s of activity. Single and multi units from both monkeys were pooled together into a pseudo population for each region as the analyses presented in this paper didn’t produce different results between the two subjects (Sup. Fig. 1 and 5). Notice that significant decoding and encoding of value and type started before or on the presentation of reward predicting cue because of the smoothing method described above.

### Unit encoding

The firing rate of units was modeled with a linear regression with value, type of reward and their interaction as variables in the epochs described above. Value was defined as a categorical variable and analyzed with ANOVAs as the firing rate profile of single neurons did not always follow a linear trend with value (Sup. Fig. 2). Individual F tests for each variable were applied on each time bin, and a bin of activity was considered to encode a variable if the p-value associated with the test was part of at least 7 consecutive time bins where the p-value was lower than 0.01. Encoding strength was represented as the negative log of the F test p-value.

For the sequential encoding analysis, trials were randomly split in half and the above encoding analysis was applied independently on each half on the delay activity, i.e. from reward predicting cue offset to reward delivery. We only kept the neurons that had at least 7 bins with an encoding p-value less than 0.01 in both training and testing sets, hence the lower number of neurons in Fig. 3.

### Decoding

Population representations of value were assessed with a ridge regression classifier (linear regression with l2 “ridge” regularization), which is a robust and efficient classifier as compared to SVM or logistic regression, which require costly computations. A one-versus-all approach was used to classify the 4 different classes of “value”, which is equivalent to 4 binary classifiers. The regularization parameter was independently optimized by exploring every power of ten between -5 and 10 with a 0.1 step for all the combinations of OFC or ACC populations, and whole population versus subspace to maximize the average performance of the classifier trained and tested on each time point between cue onset and reward. The parameter eliciting the best decoding for each condition was used for all decoding results presented here. Reported accuracies correspond to the average of a 10-fold cross-validation, to ensure generalizability of the results.

The significance of decoding accuracy was determined with permutation testing. The value label of trials was shuffled 2000 times and compared to the original data. The p-value was calculated as the number of permutations with a higher accuracy than the original data divided by 2000.

For cross-temporal decoding, we trained and tested the decoder on all possible combinations of time bin pairs. Accuracy along the diagonal represents training and testing at the same time point, while points away from the diagonal correspond to more temporally distant activities. In this case, the testing data still corresponds to out-of-sample trials so that the decoder was not trained and tested on the same trials, regardless of whether training and testing data points were from the same or different time bins.

### Subspace

To extract a stable value representation, we implemented the following method inspired by [Murray et al.2017]. The firing rate activity of each neuron in a population was averaged over time from cue onset to reward delivery, and then averaged across the trials where the cue predicted the same value. The resulting matrix had 4 rows corresponding to the 4 values and n columns corresponding to the number of neurons in the population. A principal component analysis (PCA) was performed on this matrix and allowed us to project the neural population activity of each trial at every time bin onto a 3 dimensional subspace (4 values - 1 dimension). This method finds the dimensions that capture the highest variance related to value while discarding temporal information by averaging across time.

When decoding with the subspace, we used cross-validation and ensured that the subspace was derived without the testing (out-of-sample) data. The ridge classifier was trained on the training data projected onto the subspace and then tested on the out-of-sample data projected onto the subspace obtained with the training data.

### Ensembles

Testing all the possible combinations of units to find the ensemble that best decodes value is computationally infeasible. To remedy this, [Backen et al.2018] proposed a method that finds an ensemble that substantially optimizes decoding accuracy with a subset of the full population in a computationally feasible way, though it does not guarantee the selected ensemble is the single best. This method begins by screening each unit individually and selecting the most discriminating, then successively screens the remainder and adds the unit that most increases accuracy. Effectively, this method sorts the units in the order of highest contribution to decoding accuracy. However, decoding relies on the interaction of unit activities and by successively adding units, that method does not take into account all possible interactions. For example in a 3 neuron population, neuron 2 and 3 together might yield a higher decoding accuracy than any other pair of neurons, but neuron 1 alone yields the best accuracy and might be selected first, dismissing the possibility that neuron 2 and 3 are selected as the best 2-neuron ensemble. To partially circumvent this, we used the opposite method: successively removing units. As with the adding of units, we iteratively removed the units that maximized the decoding. This method yielded better results than adding the units (Sup. Fig. 6).

To combine the ensemble and subspace approaches, this method was applied to find the ensemble that maximized the averaged cross-temporal decoding accuracy across all data points from cue onset to the reward delivery. This corresponds to averaging across all the points of the accuracy matrices represented in the figures except the points before cue and after reward. Once the best ensembles for each case (e.g. with or without subspace) were determined, cross-temporal decoding was performed with data beyond this period to show the boundaries of decoding accuracy.

### Unit and population measures

Three measures were defined to quantify the encoding of value in units. The encoding strength is the average of the negative log of the p-value across time during the delay:

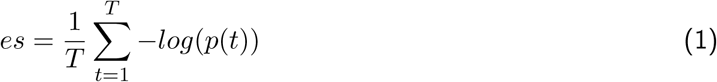

where *T* is the total number of bins in the delay, and *p*(*t*) is the p-value of a unit at time bin *t* obtained from the ANOVA F-test described in the unit encoding section above.

The encoding duration is the number of bins where the above p-value was lower than 0.01 divided by the total number of bins.

The stability measure was defined as follow:

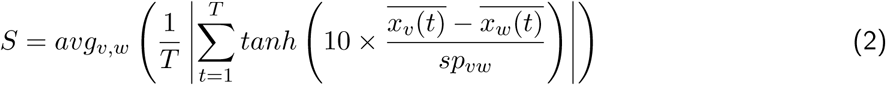

where 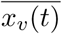 is the average firing rate of a unit for value *v, avg*_*v,w*_() is the average over all the possible combinations of values *v* and *w* except *v* = *w, tanh* is the ] — 1, 1[ bounded hyperbolic tangent function, | · | is the absolute value function and *sp*_*vw*_ is the pooled standard deviation of the firing rate of a unit across trials with value *v* and *w*:

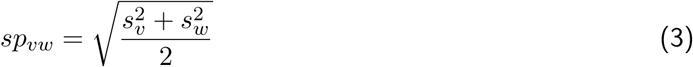

where 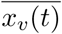 is the variance of the firing rate for trials with value *v*. Note that the difference of firing rate averaged by value, 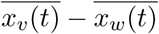, is signed and the measure takes the average of this signed difference. If a unit reverses its relative encoding of two values, the sign of the average difference changes over time and averaging across time will be close to zero, reflecting the lack of stability of encoding of that unit.

Two measures were used to quantify the contribution of units to the value subspace and the ensemble. For the value subspace we used the absolute value of the subspace weights, which corresponds to the eigenvector components obtained through a PCA of the time and trial averaged data. The contribution of a unit to the stable subspace through the ensemble method was estimated by calculating the difference in average CTD accuracy (across all time bin pairs) by removing that unit from the ensemble. The last unit removed from the ensemble was discarded. For visualization of this specific measure, a symmetric square root transformation was applied where the absolute value of negative values was used before reassigning them their sign:

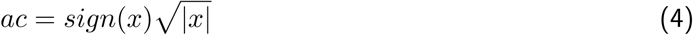

The distribution of this measure across units was asymmetrical around the zero, so correlation analyses were applied individually on positive and negative contributions (see Main text).

A “locality” measure was defined to optimize the ensemble of neurons eliciting a local CTD accuracy of value during the delay. We fit a Gaussian curve to the accuracies obtained from training at one time bin and testing on all delay bins:

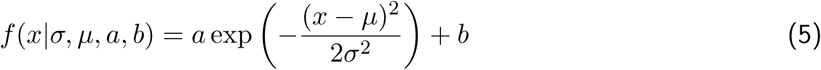

The offset value *b* was fixed to 0.25, the chance level, and the mean of the Gaussian *µ* was set to the time of the training. Only the scaling factor *a* and the standard deviation σ were optimized. The locality measure *lm* for a given ensemble is as follow:

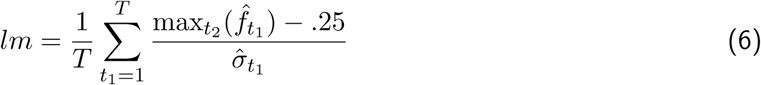

where *t*_1_ and *t*_2_ are the training time and testing time, respectively, 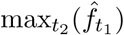 is the maximum of the Gaussian fitted on training at time *t*_1_ and testing on all *t*_2_ time bins, and 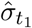 is the standard deviation of the Gaussian fitted at time *t*_1_. The contribution of a unit to the dynamic ensemble was calculated as the difference in locality measure when that unit is added to the ensemble.

### Correlations

To avoid inflated results from non-normally distributed data, we used Spearman correlations. In each of the figures showing these correlations, transformations (log or square) were applied for visualization purpose only. Note that these transformations do not affect the Spearman correlation, as it is based on rank and these transformations are monotonic and so preserve the rank of observations.

## Supplemental Data

**Supplemental Figure 1:**
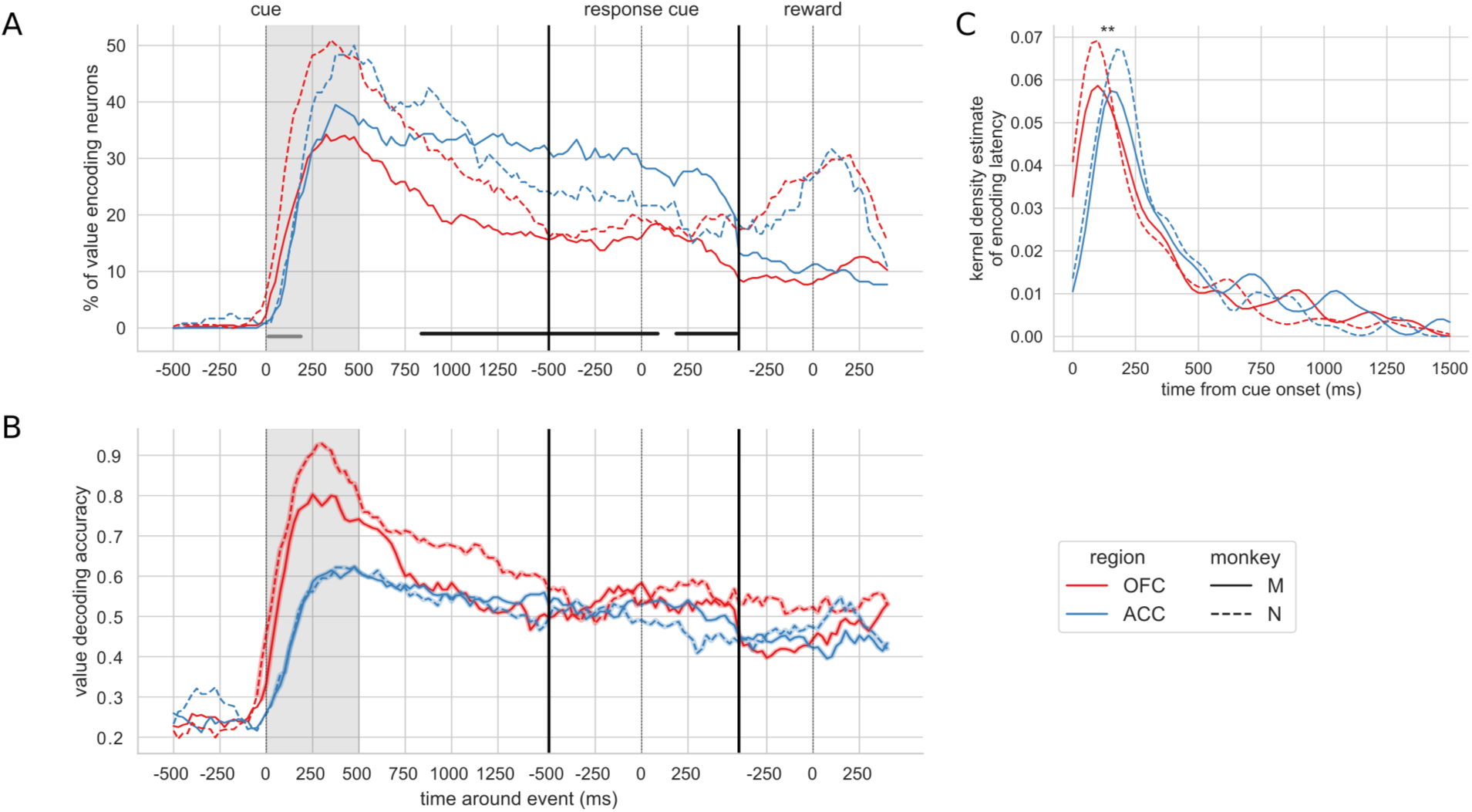
Encoding and decoding of value for each monkey. In each plot, OFC and ACC are represented with red and blue respectively, and monkey M and N with plain and dashed lines. **A** - Percentage of units encoding value at each point in time. Significant difference in value encoding neurons with Chi^2^test is indicated by thick black/grey lines at the bottom (p<0.01) for monkey M/N. **B**. - Value decoding accuracy, with a ridge regression classifier that was trained and tested independently on each time point. Significant decoding is represented by shaded area around the line and corresponds to a p-value < .01 with a 2000-permutation test. **C** - Kernel density estimate of unit value encoding latencies. OFC latencies are significantly shorter compared to ACC for both monkeys (Kruskal-Wallis, monkey M [stat=8.57, p=3.42*10^−3^] and monkey N [stat=7.74, p=5.39*10^−3^]).

**Supplemental Figure 2:**
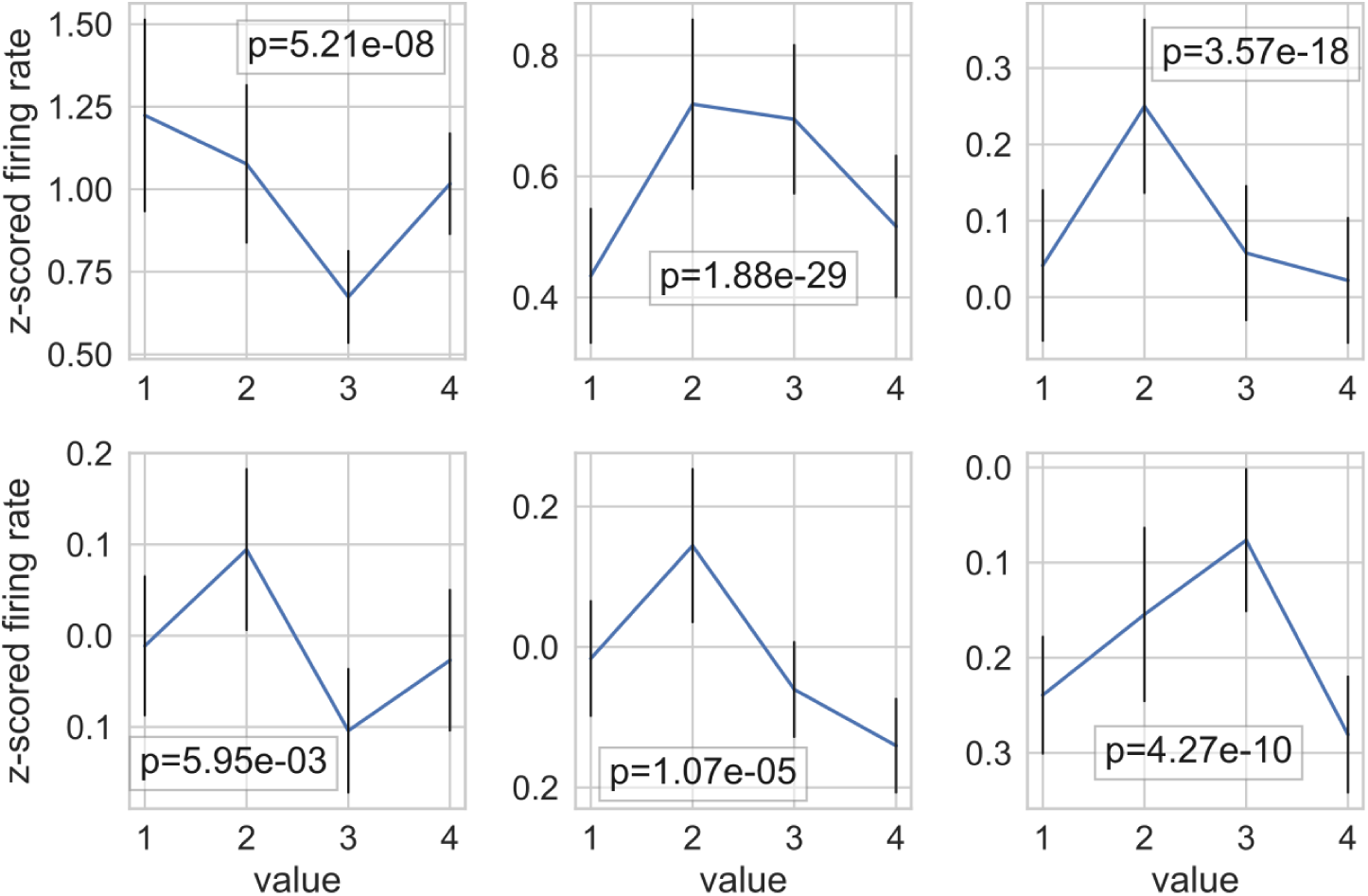
Average firing rate of 6 example neurons showing a non-linear relationship with value. Error bars represent the standard error of the mean. The p-value shown on each graph is the p-value for the factor value in an ANOVA with factors ‘type’, ‘value’ and their interaction.

**Supplemental Figure 3:**
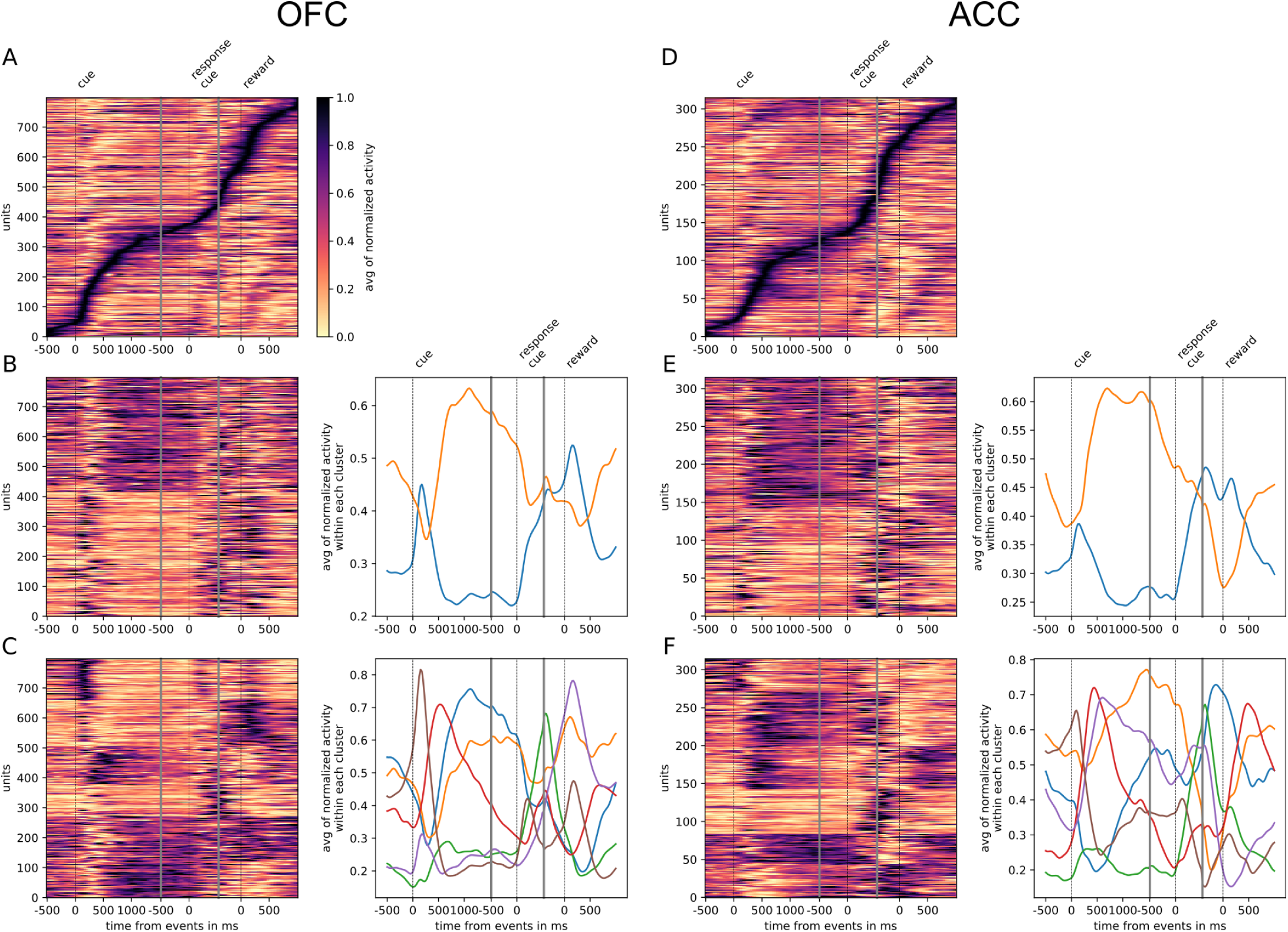
Tiling and spanning of delay with unit firing rate. Activity of neurons has been averaged across trials and normalized so that activity is scaled between 0 and 1. **A** - Tiling of delay OFC activity and surrounding epochs. **B** - Clustering of OFC activity in 2 clusters with K-means clustering. Left graph shows the activity of each unit sorted by cluster. In the right graph each curve is the averaged activity across all the units within a cluster. **C** - Same as **B** with 6 clusters (determined with elbow method) instead of 2. **D**, **E** and **F** are identical to **A**, **B** and **C** with ACC data instead of OFC. Events occurring at times indicated by 0 on the x-axis are: reward cue on, joystick instruction cue on, and reward delivery.

**Supplemental Figure 4:**
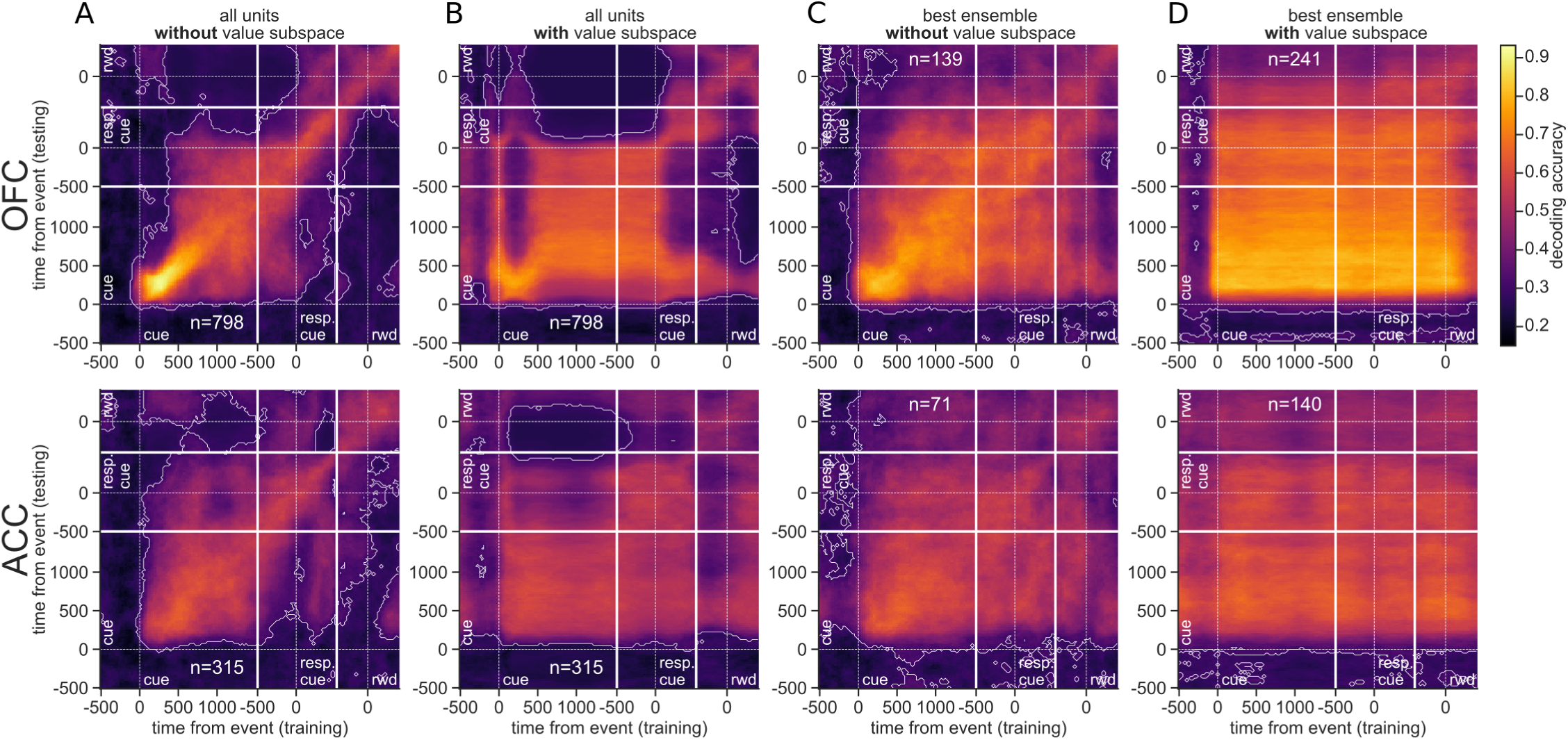
Cross-temporal decoding accuracy of value with different methods. CTD without (**A**, **C**) or with (**B**, **D**) a value subspace, and with the full population (**A**, **B**) or with the best ensemble (**C**, **D**). Thick white lines indicate junctions between the three successive epochs (cue presentation, joystick task and reward delivery). The thin dashed white lines indicate the reference event of each epoch (reward predicting cue onset, response instruction, reward). The contour curve indicates areas with a p-value lower than 0.01 with a 2000 fold permutation test. (rwd = reward, resp. cue = response cue)

**Supplemental Figure 5:**
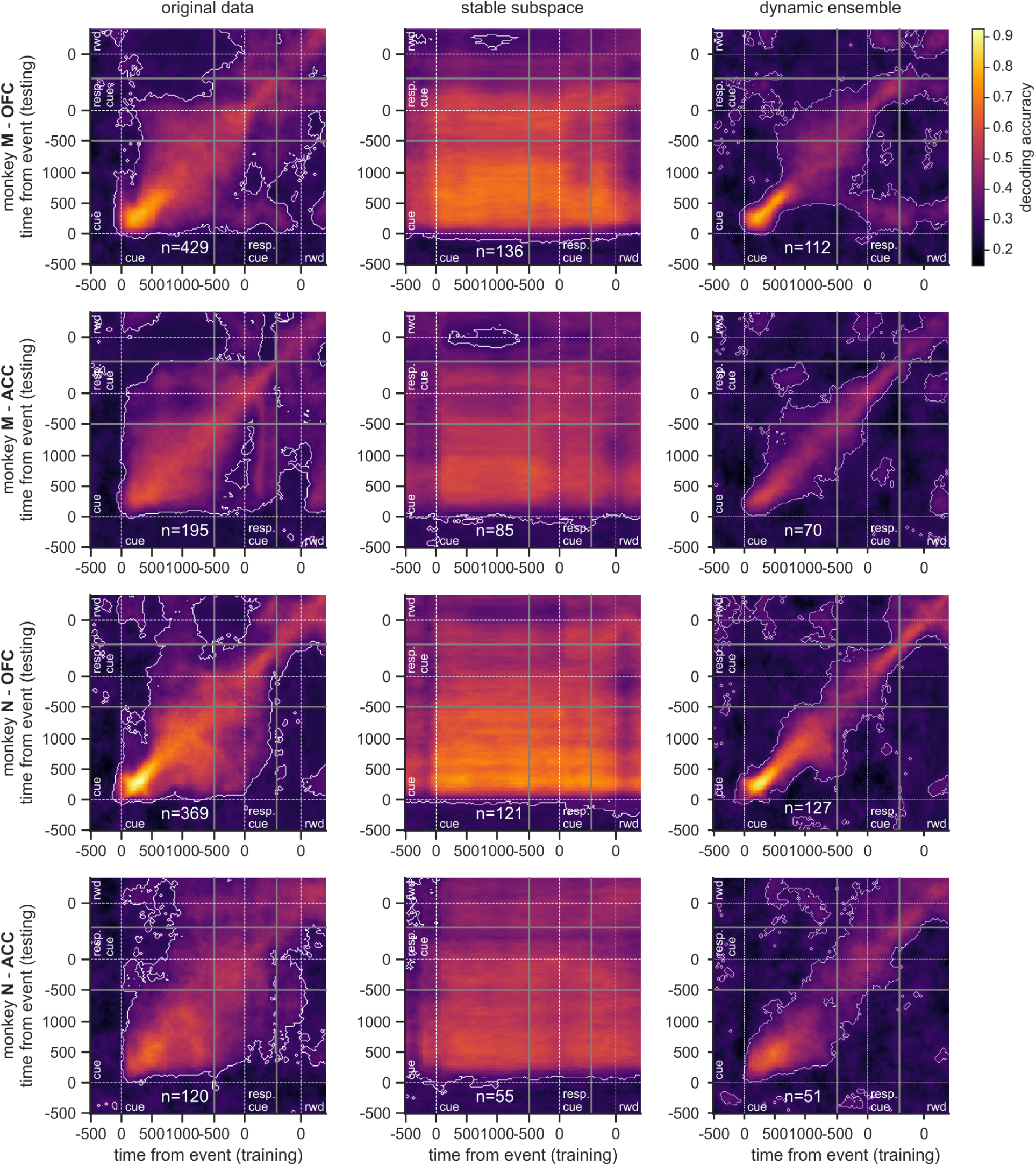
Cross-temporal decoding accuracy of value with different methods for each monkey. This figure is similar to figure 5. Each row correspond to a different combination of monkey and region, as indicated at the left of each row. Since results are similar across rows, they are largely independent of monkey and region.

**Supplemental Figure 6:**
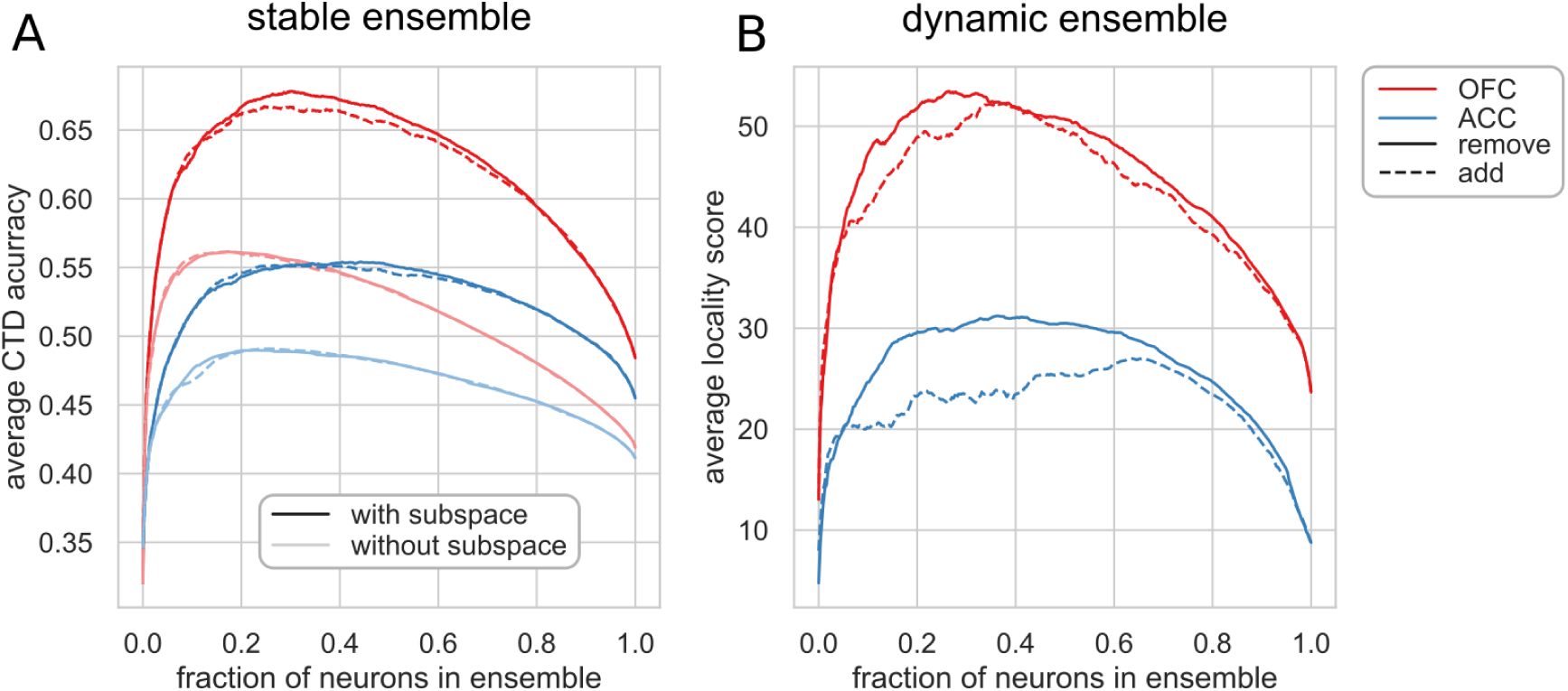
Iterative removal of units elicits better maxmimum performance than iterative addition of units. **A** - Average CTD accuracy as a function of the fraction of the total number of neurons selected to be part of the stable ensemble. **B** - Same as **A** for the dynamic ensemble with the average locality score being the maximized measure. Note the strong influence of the unit selection method (adding vs removing units) on the dynamic ensemble performances.

## References

Amiez C, Joseph JP, Procyk E (2006) Reward encoding in the monkey anterior cingulate cortex. Cereb. cortex 16:1040–55.

Amit DJ (1995) The Hebbian paradigm reintegrated: Local reverberations as internal representations. Behav. brain siences 18:617–657.

Asaad WF, Eskandar EN (2008) A flexible software tool for temporally-precise behavioral control in Matlab. J. Neurosci. Methods 174:245–258.

Astrand E, Enel P, Ibos G, Dominey PF, Baraduc P, Ben Hamed S (2014) Comparison of classifiers for decoding sensory and cognitive information from prefrontal neuronal populations. PLoS One 9:e86314.

Backen T, Treue S, Martinez-Trujillo JC (2018) Encoding of Spatial Attention by Primate Prefrontal Cortex Neuronal Ensembles. Eneuro 5:ENEURO.0372–16.2017.

Barak O, Rigotti M, Fusi S (2013) The sparseness of mixed selectivity neurons controls the generalization-discrimination trade-off. J. Neurosci. 33:3844–56.

Barak O, Tsodyks M, Romo R (2010) Neuronal population coding of parametric working memory. J. Neurosci. 30:9424–30.

Cavanagh SE, Towers JP, Wallis JD, Hunt LT, Kennerley SW (2018) Reconciling persistent and dynamic hypotheses of working memory coding in prefrontal cortex. Nat. Commun. 9:3498.

Compte A, Brunel N, Goldman-Rakic PS, Wang XJ (2000) Synaptic Mechanisms and Network Dynamics Underlying Spatial Working Memory in a Cortical Network Model. Cereb. Cortex 10:910–923.

Constantinidis C, Funahashi S, Lee D, Murray JD, Qi XL, Wang M, Arnsten AF (2018) Persistent Spiking Activity Underlies Working Memory. J. Neurosci. 38:7020–7028.

Constantinidis C, Klingberg T (2016) The neuroscience of working memory capacity and training. Nat. Rev. Neurosci. 17:438–449.

Cueva CJ, Marcos E, Saez A, Genovesio A, Jazayeri M, Romo R, Salzman CD, Shadlen MN, Fusi S (2019) Low dimensional dynamics for working memory and time encoding. bioRxiv p. 504936.

Eichenbaum HB (2014) Time cells in the hippocampus: a new dimension for mapping memories. Nat. Rev. Neurosci. 15.

Enel P, Procyk E, Quilodran R, Dominey PF (2016) Reservoir Computing Properties of Neural Dynamics in Prefrontal Cortex. PLOS Comput. Biol. 12:e1004967.

Fujisawa S, Amarasingham A, Harrison MT, Buzsáki G (2008) Behavior-dependent short-term assembly dynamics in the medial prefrontal cortex. Nat. Neurosci. 11:823–833.

Funahashi S (1989) Mnemonic coding of visual space in the monkey’s dorsolateral prefrontal cortex. J. Neurosci. 6:331–349.

Fuster JM (1973) Unit activity in prefrontal cortex during delayed-response performance: neuronal correlates of transient memory. J. Neurophysiol. 36:61–78.

Goldman-Rakic PS (1996) Regional and cellular fractionation of working memory. Proc. Natl. Acad. Sci. 93:13473–13480.

Harvey CD, Coen P, Tank DW (2012) Choice-specific sequences in parietal cortex during a virtual-navigation decision task. Nature 484:62–68.

Jacob SN, Nieder A (2014) Complementary roles for primate frontal and parietal cortex in guarding working memory from distractor stimuli. Neuron 83:226–237.

Jaeger H (2001) The” echo state” approach to analysing and training recurrent neural networks Technical report, Technical Report GMD Report 148, German National Research Center for Information Technology.

Kennerley SW, Dahmubed AF, Lara AH, Wallis JD (2009) Neurons in the frontal lobe encode the value of multiple decision variables. J. Cogn. Neurosci. 21:1162–78.

Kennerley SW, Wallis JD (2009) Encoding of reward and space during a working memory task in the orbitofrontal cortex and anterior cingulate sulcus. J. Neurophysiol. 102:3352–64.

Lara AH, Kennerley SW, Wallis JD (2009) Encoding of Gustatory Working Memory by Orbitofrontal Neurons. J. Neurosci. 29:765–774.

Lebedev MA, Messinger A, Kralik JD, Wise SP (2004) Representation of Attended Versus Remembered Locations in Prefrontal Cortex. PLoS Biol. 2:e365.

Lundqvist M, Herman P, Miller EK (2018) Working Memory: Delay Activity, Yes! Persistent Activity? Maybe Not. J. Neurosci. 38:7013–7019.

Lundqvist M, Rose J, Herman P, Brincat S, Buschman T, Miller EK (2016) Gamma and Beta Bursts Underlie Working Memory. Neuron 90:152–164.

Maass W, Joshi P, Sontag ED (2007) Computational aspects of feedback in neural circuits. PLoS Comput. Biol. 3:e165.

Maass W, Natschläger T, Markram H (2002) Real-time computing without stable states: a new framework for neural computation based on perturbations. Neural Comput. 14:2531–60.

MacDonald CJ, Lepage KQ, Eden UT, Eichenbaum HB (2011) Hippocampal “time cells” bridge the gap in memory for discontiguous events. Neuron 71:737–749.

Machens CK, Romo R, Brody CD (2005) Flexible Control of Mutual Inhibition: A Neural Model of Two-Interval Discrimination. Science (80-.). 307:1121–4.

Masse NY, Yang GR, Song HF, Wang XJ, Freedman DJ (2019) Circuit mechanisms for the maintenance and manipulation of information in working memory. Nat. Neurosci. .

Meyers EM (2018) Dynamic population coding and its relationship to working memory. J. Neurophysiol. 120:2260–2268.

Meyers EM, Freedman DJ, Kreiman G, Miller EK, Poggio T (2008) Dynamic population coding of category information in inferior temporal and prefrontal cortex. J. Neurophysiol. 100:1407–19.

Miller KJ, Botvinick MM, Brody CD (2018) Value Representations in Orbitofrontal Cortex Drive Learning, but not Choice. bioRxiv pp. 1–25.

Murray JD, Bernacchia A, Roy NA, Constantinidis C, Romo R, Wang XJ (2017) Stable population coding for working memory coexists with heterogeneous neural dynamics in prefrontal cortex. Proc. Natl. Acad. Sci. 114:394–399.

Orhan AE, Ma WJ (2019) A diverse range of factors affect the nature of neural representations underlying short-term memory. Nat. Neurosci. 22:275–283.

Padoa-Schioppa C (2011) Neurobiology of Economic Choice: A Good-Based Model. Annu. Rev. Neurosci. 34:333–359.

Parthasarathy A, Herikstad R, Bong JH, Medina FS, Libedinsky C, Yen SC (2017) Mixed selectivity morphs population codes in prefrontal cortex. Nat. Neurosci. 20:1770–1779.

Pascanu R, Jaeger H (2011) A neurodynamical model for working memory. Neural Networks 24:199–207.

Pastalkova E, Itskov V, Amarasingham A, Buzsaki G (2008) Internally Generated Cell Assembly Sequences in the Rat Hippocampus. Science (80-.). 321:1322–1327.

Rainer G, Miller EK (2002) Timecourse of object-related neural activity in the primate prefrontal cortex during a short-term memory task. Eur. J. Neurosci. 15:1244–1254.

Rajan K, Harvey CD, Tank DW (2015) Recurrent Network Models of Sequence Generation and Memory. Neuron 90:1–15.

Rich EL, Wallis JD (2016) Decoding subjective decisions from orbitofrontal cortex. Nat. Neurosci. 19:973–980.

Rich EL, Wallis JD (2017) Spatiotemporal dynamics of information encoding revealed in orbitofrontal high-gamma. Nat. Commun. 8:1–13.

Riley MR, Constantinidis C (2016) Role of Prefrontal Persistent Activity in Working Memory. Front. Syst. Neurosci. 9:1–14.

Rushworth MFS, Behrens TEJ (2008) Choice, uncertainty and value in prefrontal and cingulate cortex. Nat. Neurosci. 11:389–397.

Seung HS (1996) How the brain keeps the eyes still. Proc. Natl. Acad. Sci. 93:13339–13344.

Spaak E, Watanabe K, Funahashi S, Stokes MG (2017) Stable and Dynamic Coding for Working Memory in Primate Prefrontal Cortex. J. Neurosci. 37:6503–6516.

Stokes MG, Buschman TJ, Miller EK (2017) Dynamic Coding for Flexible Cognitive Control In Egner T, editor, Wiley Handb. Cogn. Control, pp. 221–241. John Wiley & Sons, Ltd, Chichester, UK.

Stokes MG, Kusunoki M, Sigala N, Nili H, Gaffan D, Duncan J (2013) Dynamic Coding for Cognitive Control in Prefrontal Cortex. Neuron 78:1–12.

Stoll FM, Fontanier V, Procyk E (2016) Specific frontal neural dynamics contribute to decisions to check. Nat. Commun. 7:11990.

Wang XJ (2001) Synaptic reverberation underlying mnemonic persistent activity. Trends Neurosci. 24:455–63.

Warden MR, Miller EK (2007) The Representation of Multiple Objects in Prefrontal Neuronal Delay Activity. Cereb. Cortex 17:i41–i50.

Wasmuht DF, Spaak E, Buschman TJ, Miller EK, Stokes MG (2018) Intrinsic neuronal dynamics predict distinct functional roles during working memory. Nat. Commun. 9.

Watanabe K, Funahashi S (2014) Neural mechanisms of dual-task interference and cognitive capacity limitation in the prefrontal cortex. Nat. Neurosci. 17:601–611.

